# Are skyline plot-based demographic estimates overly dependent on smoothing prior assumptions?

**DOI:** 10.1101/2020.01.27.920215

**Authors:** Kris V Parag, Oliver G Pybus, Chieh-Hsi Wu

## Abstract

In Bayesian phylogenetics, the coalescent process provides an informative framework for inferring changes in the effective size of a population from a phylogeny (or tree) of sequences sampled from that population. Popular coalescent inference approaches such as the *Bayesian Skyline Plot, Skyride* and *Skygrid* all model these population size changes with a discontinuous, piecewise-constant function but then apply a smoothing prior to ensure that their posterior population size estimates transition gradually with time. These prior distributions implicitly encode extra population size information that is not available from the observed coalescent data i.e., the tree. Here we present a novel statistic, Ω, to quantify and disaggregate the relative contributions of the coalescent data and prior assumptions to the resulting posterior estimate precision. Our statistic also measures the additional mutual information introduced by such priors. Using Ω we show that, because it is surprisingly easy to over-parametrise piecewise-constant population models, common smoothing priors can lead to overconfident and potentially misleading inference, even under robust experimental designs. We propose Ω as a useful tool for detecting when effective population size estimates are overly reliant on prior assumptions and for improving quantification of the uncertainty in those estimates.

## Introduction

The coalescent process models how changes in the effective size of a target population influence the phylogenetic patterns of sequences sampled from that population. First derived in [16] under the assumption of a constant sized population, the coalescent process has since been extended to account for temporal variation in the population size [12], structured demographics [1] and multi-locus sampling [19]. Inference under these models aims to statistically recover the unknown effective population size (or demographic) history from the reconstructed phylogeny (or tree) and has provided insights into infectious disease epidemiology, population genetics and molecular ecology [31, 38, 25]. Here we focus on coalescent processes that describe the genealogies of serially-sampled individuals from populations with deterministically varying size. These are widely applied to study the phylodynamics of infectious diseases [12, 29].

Early approaches to inferring effective population size from coalescent phylogenies used pre-defined parametric models (e.g. exponential or logistic growth functions) to represent temporal demographic changes [17, 25]. While these formulations required only a few variables and provided interpretable estimates, selecting the most appropriate parametric description could be challenging and risk underfitting complex trends [20]. This motivated the introduction of the *classic skyline plot* [26], which, by proposing an independent, piecewise-constant demographic change at every coalescent event (i.e at branching times in the phylogeny), maximised flexibility and removed parametric restrictions. However, this flexibility came at the cost of increased estimation noise and potential overfitting of changes in effective population size [13].

Efforts to redress these issues within a piecewise-constant framework subsequently spawned a family of skyline plot-based methods [13]. Among these, the most popular and commonly-used are the *Bayesian Skyline Plot* (BSP) [8], the *Skyride* [20] and the *Skygrid* [11] approaches. All three attempted to regulate the sharp fluctuations of the inferred piecewise-constant demographic function by enforcing *a priori* assumptions about the smoothness (i.e. the level of autocorrelation among piecewise-constant segments) of real population dynamics. This was seen as a biologically sensible compromise between noise regulation and model flexibility [21, 34].

The BSP limited overfitting by (i) predefining fewer piecewise demographic changes than coalescent events and (ii) smoothing noise by asserting *a priori* that the population size after a change-point was exponentially distributed around the population size before it. This method was questioned by [20] for making strong smoothing and change-point assumptions and stimulated the development of the Skyride, which embeds the flexible classic skyline plot within a tunable Gaussian smoothing field. The Skygrid, which extends the Skyride to multiple loci and allows arbitrary change-points (the BSP and Skyride change-times coincide with coalescent events), also uses this prior. The Skyride and Skygrid methods aimed to better trade off prior influence with noise reduction, and while somewhat effective, are still imperfect because they can fail to recover genuinely abrupt demographic changes such as bottlenecks [9].

As a result, studies continue to explore and address the non-trivial problem of optimising this tradeoff, either by searching for less-restrictive and more adaptive priors [9] or by deriving new data-driven skyline change-point grouping strategies [21]. The evolution of coalescent model inference thus reflects a desire to understand and fine-tune how prior assumptions and observed phylogenetic data interact to yield reliable posterior population size estimates. Surprisingly, and in contrast to this desire, no study has yet tried to directly and rigorously measure the relative influence of the priors and data on these estimates.

Here we develop and present a novel information theoretic statistic, Ω, to formally quantify and disaggregate the contributions of both priors and data on the uncertainty around the posterior demographic estimates of popular skyline-based coalescent methods. Using Ω we show how widely-used smoothing priors can result in overconfident population size inferences (i.e. estimates with unjustifiably small credible intervals) and provide practical guidelines against such circumstances. We illustrate the utility of this approach on well-characterised datasets describing the population size of HCV in Egypt [25] and ancient Beringian steppe Bison [31].

To our knowledge, Ω, which in theory can be adapted to any prior-data comparison problem, is new not only to the field of phylogenetics but also across statistics and data science. While inference that is strongly driven by prior assumptions can be beneficial, for example when a prior encodes expert knowledge or salient dynamics, having a measure of the relative information introduced by data and prior distributions can improve the reproducibility and interpretability of analyses. Our statistic will help to detect when prior assumptions are inadvertently and overly influencing demographic estimates and will hopefully serve as a diagnostic tool that future methods can employ to optimise and validate their prior-data tradeoffs.

## I. Materials and Methods

### A. Coalescent Inference

We provide an overview of the coalescent process and statistical inference under skyline plot-based demographic models. The coalescent is a stochastic process that describes the ancestral genealogy of sampled individuals or lineages from a target population [16]. Under the coalescent, a tree or phylogeny of relationships among these individuals is reconstructed backwards in time with coalescent events defined as the points where pairs of lineages merge (i.e. coalesce) into their ancestral lineage. This tree, 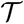, is rooted at time *T* into the past, which is the time to the most recent common ancestor (TMRCA) of the sample. The tips of 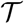 correspond to sampled individuals.

The rate at which coalescent events occur (i.e. the rate of branching in 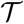) is determined by and hence informative about the effective size of the target population. We assume that a total of *n* ≥ 2 samples are taken from the target population at *n_s_* ≥ 1 distinct sampling times, which are independent of and uninformative about population size changes [8]. We do not specify the sample generating process as it does not affect our analysis by this independence assumption [24]. We let *c_i_* be the time of the *i*^th^ coalescent event in 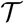 with 1 ≤ *i* ≤ *n* – 1 and *c*_*n*–1_ = *T* (*n* samples can coalesce *n* – 1 times before reaching the TMRCA).

We use *l_t_* to count the number of lineages in 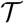 at time *t* ≥ 0 into the past; *l_t_* then decrements by 1 at every *c_i_* and increases at sampling times. Here *t* = 0 is the present. The effective population size or demographic function at *t* is *N*(*t*) so that the coalescent rate underlying 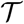 is 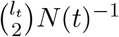 [16]. While *N*(*t*) can be described using appropriate parametric formulations [23], it is more common to represent *N*(*t*) by some tractable *p*-dimensional piecewise-constant approximation [13]. Thus, we can write 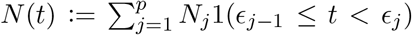, with *p* ≥ 1 as the number of piecewise-constant segments. Here *N_j_* is the constant population size of the *j*^th^ segment which is delimited by times [*ϵ*_*j*–1_, *ϵ_j_*), with *ϵ*_0_ = 0 and *ϵ_p_* ≥ *T* and 1(*x*) is an indicator function. The rate of producing new coalescent events is then 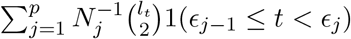. Kingman’s coalescent model is obtained by setting *p* = 1 (constant population of *N*_1_).

When reconstructing the population size history of infectious diseases, it is often of interest to infer *N*(*t*) from 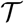 [13], which forms our coalescent data generating process. If ***N*** = [*N*_1_,…, *N_p_*] denotes the vector of demographic parameters to be estimated then the coalescent data log-likelihood 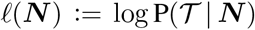 can be obtained from [24] [33] as

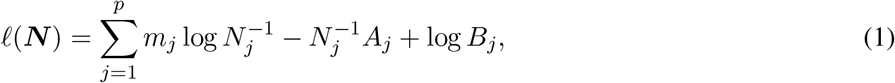

with *A_j_* and *B_j_* as constants that depend on the times and lineage counts of the *m_j_* coalescent events that fall within the *j*^th^ segment duration [*ϵ*_*j*–1_, *ϵ_j_*), and 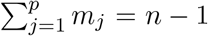. Eq. (1) is equivalent to the standard serially-sampled skyline log-likelihood in [8], except that we do not restrict *N*(*t*) to change only at coalescent event times.

In Bayesian phylogenetic inference, skyline-based methods such as the BSP, Skyride and Skygrid combine this likelihood with a prior distribution P(***N***), which encodes *a priori* beliefs about the demographic function. This yields a population size posterior, from Bayes law, which depends on both the prior and coalescent data-likelihood as:

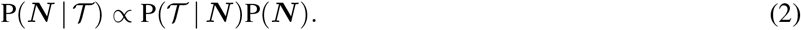

Here we assume that the phylogeny, 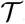, is known without error. In some instances, only sampled sequence data, ***D***, are available and a distribution over 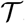 must be reconstructed from ***D*** under a model of molecular evolution with parameters ***θ***. Eq. (2) is then embedded in the more complex expression 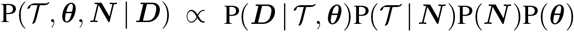, which involves inferring both the tree and population size [7].

While we do not consider this extension here we note that results presented here are still applicable and relevant. This follows because the output of the more complex Bayesian analysis above (i.e. when sequence data ***D*** are used directly) is a posterior distribution over tree space. We can sample from this posterior and treat each sampled tree effectively as a fixed tree. Consequently, we expect any summary statistic that we derive here, under the assumption of a fixed-tree will be usable in studies that incorporate genealogical uncertainty by computing the distribution of that statistic over this covering set of sampled posterior trees.

### B. Information and Estimation Theory

We review and extend some concepts from information and estimation theory as applied to skyline-based coalescent inference. We consider a general parametrisation of the effective population size ***ψ*** = [*ψ*_1_,…, *ψ_p_*], where *ψ_i_* = *ϕ*(*N_i_*) for all *i* ∈ {1,…, *p*} and *ϕ*(·) is a differentiable function. Popular skyline-based methods usually choose the identity function (e.g. BSP) or the natural logarithm (e.g. the Skyride and Skygrid) for *ϕ*. Eq. (1) and Eq. (2) are then reformulated with 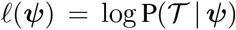 as the coalescent data log-likelihood and P(***ψ***) as the demographic prior. The Bayesian posterior, 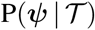 combines this likelihood and prior, and hence is influenced by both the coalescent data and prior beliefs. We can formalise these influences using information theory.

The expected Fisher information, 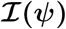, is a *p* × *p* matrix with (*i*, *j*)^th^ element 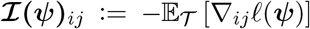 [18]. The expectation is taken over the coalescent tree branches and 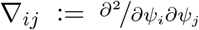. As observed in [24], 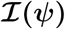 quantifies how precisely we can estimate the demographic parameters, ***ψ***, from the coalescent data, 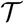. Precision is defined as the inverse of variance [18]. The BSP, Skyride and Skygrid parametrisations all yield 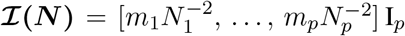 and 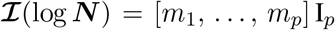, with **I***_p_* as a *p* × *p* identity matrix [24]. These matrices provide several useful insights that we will exploit in later sections. First, 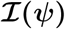 is orthogonal (diagonal), meaning that the coalescent process over the *j*^th^ segment [*ϵ*_*j*–1_, *ϵ_j_*) can be treated as deriving from an independent Kingman coalescent with constant population size *N_j_* [23]. Second, the number of coalescent events in that segment, *m_j_*, controls the Fisher information available about *N_j_*. Last, working under log *N_j_* removes any dependence of this Fisher information component on the unknown parameter *N_j_* [24].

The prior distribution, P(***ψ***), that is placed on the demographic parameters can alter and impact both estimate bias and precision. We can gauge prior-induced bias by comparing the maximum likelihood estimate (MLE), 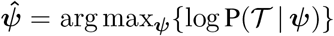 with the maximum a posteriori estimate (MAP), 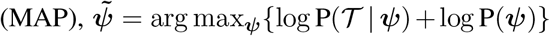 [36]. The difference 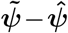 measures this bias. We can account for prior-induced precision by computing Fisher-type matrices for the prior and posterior as 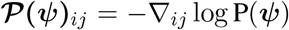 and 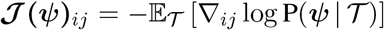 [35, 14]. Combining these gives

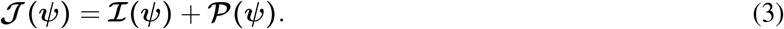

Eq. (3) shows how the posterior Fisher information matrix, 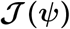, relates to the standard Fisher information 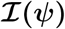 and the prior second derivative 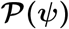. We make the common regularity assumptions (see [14] for details) that ensure 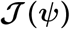 is positive definite and that all Fisher matrices exist. These assumptions are valid for exponential families such as the piecewise-constant coalescent [18, 24]. Eq. (3) will prove fundamental to resolving the relative impact of the prior and data on the best precision achievable using 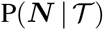. We also define expectations on these matrices with respect to the prior as 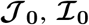 and 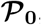, with 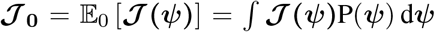, for example. These matrices are now constants instead of functions of ***ψ***. Eq. (3) also holds for these matrices [35].

These Fisher information matrices set theoretical upper bounds on the precision attainable by all possible statistical inference methods. For any unbiased estimate of 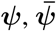 the Cramer-Rao bound (CRB) states that 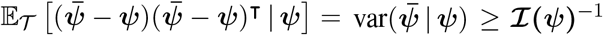 with 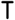 indicating transpose. If we relax the unbiased requirement and include prior (distribution) information then the Bayesian or posterior Cramer-Rao lower bound (BCRB) controls the best estimate precision [36]. If 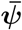 is any estimator of ***ψ*** then the BCRB states that 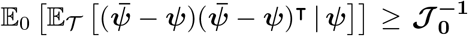. This bound is not dependent on ***ψ*** due to the extra expectation over the prior [35].

The CRB describes how precisely we can estimate demographic parameters using just the coalescent data and is achieved (asymptotically) with equality for skyline (piecewise-constant) coalescent models [24]. The BCRB, instead, defines the precision limit for the combined contributions of the data and the prior. The CRB is a frequentist bound that assumes a true fixed ***ψ***, while the BCRB is a Bayesian bound that treats ***ψ*** as a random parameter. The expectation over the prior connects the two formalisms [2]. Given their importance in delimiting precision, the 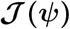 and 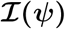 Fisher matrices will be central to our analysis, which focuses on resolving the individual contributions of the data versus prior assumptions.

## II. Results

### A. *The Coalescent Information Ratio*, Ω

We propose and derive the coalescent information ratio, Ω, as a statistic for evaluating the relative contributions of the prior and coalescent data to the posterior estimates obtained as solutions to Bayesian skyline inference problems (see Materials and Methods). Consider such a problem in which the *n*-tip phylogeny 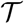 is used to estimate the *p*-element demographic parameter vector ***ψ***. Let 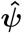 be the MLE of ***ψ*** given the coalescent data 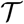. Asymptotically, the uncertainty around this MLE can be described with a multivariate Gaussian distribution with covariance matrix 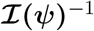. The Fisher information, 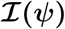 then defines a confidence ellipsoid that circumscribes the total uncertainty from this distribution. In [24] this ellipsoid was found central to understanding the statistical properties of skyline-based estimates.

The volume of this ellipsoid is 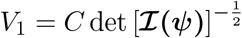, with *C* as a *p*-dependent constant. Decreasing *V*_1_ increases the best estimate precision attainable from the data 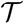 [18]. In a Bayesian framework, the asymptotic posterior distribution of ***ψ*** also follows a multivariate Gaussian distribution with covariance matrix of 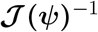. We can therefore construct an analogous ellipsoid from 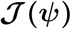 with volume 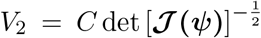 that measures the uncertainty around the MAP estimate 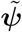 [35]. This volume includes the effect of both prior and data on estimate precision. Accordingly, we propose the ratio

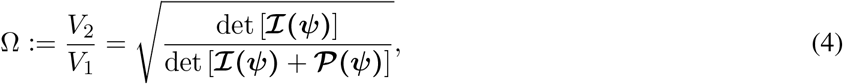

as a novel and natural statistic for dissecting the relative impact of the data and prior on posterior estimate precision.

From Eq. (4), we observe that 0 ≤ Ω ≤ 1 with Ω = 1 signifying that the information from our prior distribution is negligible in comparison to that from the data and Ω = 0 indicating the converse. Importantly, we find

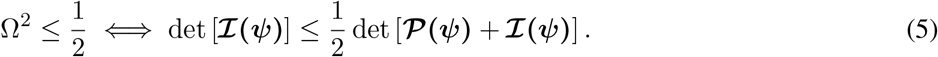

At this threshold value 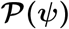 contributes at least as much information as the data. Moreover, lim_*n*→∞_ Ω = 1 since the prior contribution becomes negligible with increasing data and Ω is undefined when ***ψ*** is unidentifiable from 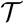 (i.e. when 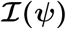 is singular [30]). Consequently, we posit that a smaller Ω implies the prior provides a greater contribution to estimate precision.

We define Ω as an information ratio due to its close connection to both the Fisher and mutual information. The mutual information between ***ψ*** and 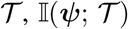, measures how much information (in bits for example) 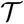 contains about ***ψ*** [6]. This is distinct but related to 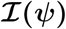, which quantifies the precision of estimating ***ψ*** from 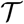 [5]. Recent work from [14] into the connection between the Fisher and mutual information has yielded two key approximations to 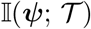. These can be obtained by substituting either 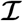 or 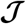 for 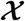 in

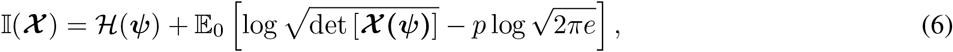

with 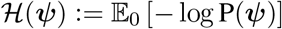 as the differential entropy of ***ψ*** [6].

For a flat prior or many observations, 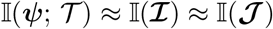, as the prior contributes little or no information [5]. For sharper priors, 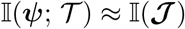 as the prior contribution is significant – using 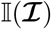 would lead to large errors [14]. Eq. (6) is predicated on (i) regularity assumptions for the distributions used (i.e. that the second derivatives exist), (ii) conditional dependence of the observed data given ***ψ*** and (iii) that the likelihood is peaked around its most probable value [18, 5, 14]. The skyline-based inference problems that we consider here automatically satisfy (i) and (ii) as these models belong to an exponential family. Condition (iii) is satisfied for moderate to large trees (and asymptotically) [18, 24].

Using the above approximations, we derive the interesting expression

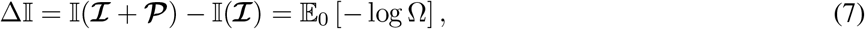

which suggests that our ratio directly measures the excess mutual information introduced by the prior, providing a substantive link between how sharper estimate precision is attained with extra mutual information. Observe that both sides of Eq. (7) diminish when 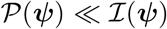. Because the mutual information and its approximations (see Eq. (6)) are invariant to invertible parameter transformations [14], our coalescent information ratio does not depend on whether we infer ***N***, its inverse, or its logarithm.

Moreover, we can use normalising transformations to make Ω valid at even small tree sizes. In [32] several such transformations for exponentially distributed models like the coalescent are derived. Among them, the log transform can achieve approximately normal log-likelihoods for about 7 observations and above (*n* ≥ 8). Thus, log ***N***, which is also optimal for experimental design [24], ensures the validity of Ω on small trees. This is the parametrisation adopted by the Skyride and Skygrid methods [20]. Other (cubic-root) parametrisations under which Ω would be valid at even smaller *n* also exist [32].

Eq. (4)–Eq. (7) are not restricted to coalescent inference problems and are generally applicable to statistical models that involve exponential families [18]. We now specify Ω for skyline-based models, which all possess piecewise-constant population sizes and orthogonal 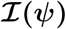 matrices [24]. These properties permit the expansion [15]:

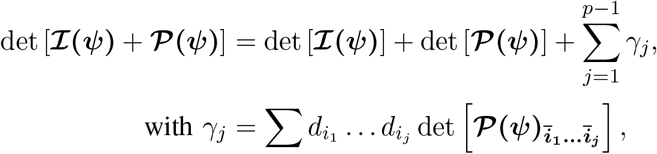

where *d_k_* are the diagonal elements of 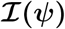 with 1 ≤ *i*_1_ <… < *j_j_* ≤ *p*, and 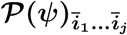 is the sub-matrix formed by deleting the (*i*_1_,…, *i_j_*)^th^ rows and columns of 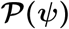.

This allows us to formulate a prior signal-to-noise ratio

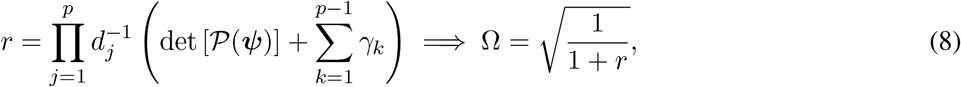

which quantifies the relative excess Fisher information (the ‘signal’) that is introduced by the prior. This ratio signifies when the prior contribution overwhelms that of the data i.e. 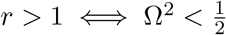. Having derived theoretically meaningful metrics for resolving prior-data precision contributions, we next investigate their ramifications.

### B. The Kingman Conjugate Prior

Kingman’s coalescent process [16], which describes the phylogeny of a constant sized population *N*_1_, is the foundation of all skyline model formulations. Specifically, a *p*-dimensional skyline model is analogous to having *p* Kingman coalescent models, the *j*^th^ of which is valid over [*ϵ*_*j*–1_, *ϵ_j_*) and describes the genealogy under population size *N_j_*. Here we use Kingman’s coalescent to validate and clarify the utility of Ω as a measure of relative data-prior precision contributions.

We assume an *n*-tip Kingman coalescent tree, 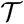 and initially work with the inverse parametrisation, 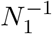. We scale 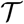 at *t* by 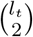 as in [23] so that 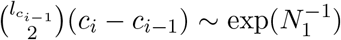 for 1 ≤ *i* ≤ *n* – 1 with *c*_0_ = 0. If *y* defines the space of 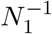 values, and has prior distribution P(*y*), then, by [33], its posterior is

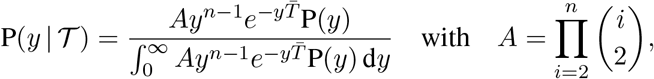

where *A* is a constant and 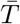 is the scaled TMRCA of 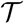.

The likelihood function embedded within 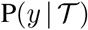 is proportional to a shape-rate parametrised gamma distribution, with known shape *n*. The conjugate prior for 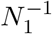 is also gamma [10] i.e. 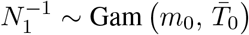 with shape *m*_0_ and rate 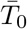. The posterior distribution is then 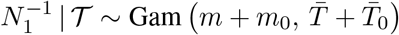 with *m* = *n* – 1 counting coalescent events in 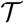 [28]. Transforming to *N*_1_ implies 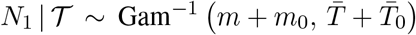. This is an inverse gamma distribution with mean 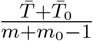, shape *m* + *m*_0_ and inverse rate 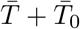. If *x* describes the space of possible *N*_1_ values and 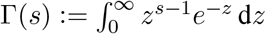 then

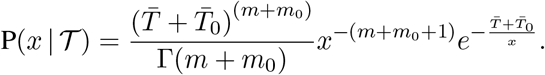

We can interpret the parameters of the gamma posterior distribution as involving a prior contribution of *m*_0_ – 1 coalescent events from a virtual tree, 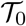, with scaled TMRCA 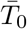. This is then combined with the actual coalescent data, which contributes *m* coalescent events from 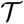, with scaled TMRCA of 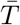 [28]. This offers a very clear breakdown of how our posterior estimate precision is derived from prior and likelihood contributions, and suggests that if 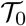 has more tips than 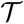 then we are depending more on the prior than the data. We now calculate Ω to determine if we can formalise this intuition.

The Fisher information values of 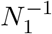 are 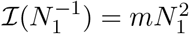 and 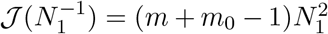. The information ratio and mutual information difference, 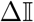, which hold for all parametrisations, then follow from Eq. (4), Eq. (7) and Eq. (8) as

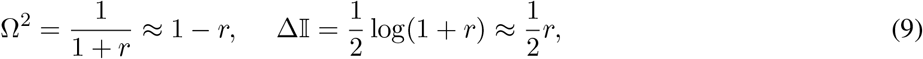

with 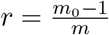, as the signal-to-noise ratio. The approximations shown are valid when *r* ≪ 1. Interestingly, when *m*_0_ – 1 = *m* so that *r* = 1, we get Ω^2^ = 1/2 (see Eq. (5)). This exactly quantifies the relative impact of real and virtual observations described previously. At this point we are being equally informed by both the conjugate prior and the likelihood. Prior over-reliance can be defined by the threshold condition of *r* > 1 ⇒ Ω^2^ < 1/2.

The expression of 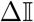 confirms our interpretation of *r* as an effective signal-to-noise ratio controlling the extra mutual information introduced by the conjugate prior. This can be seen by comparison with the standard Shannon mutual information expressions from information theory [6]. At small *r*, where the data dominates, we find that the prior linearly detracts from Ω^2^ and linearly increases 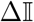. We also observe that 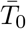, the gamma rate parameter, has no effect on estimate precision or mutual information.

Our ratio Ω therefore provides a systematic decomposition of the posterior population size estimate precision and generalises the virtual observation idea to any prior distribution. In essence, the prior is contributing an effective sample size, which for the conjugate Kingman prior is *m*_0_ – 1. We summarise these points in Fig. 1, which shows the conjugate prior and two posteriors together with their corresponding Ω^2^ values.

**Fig. 1:**
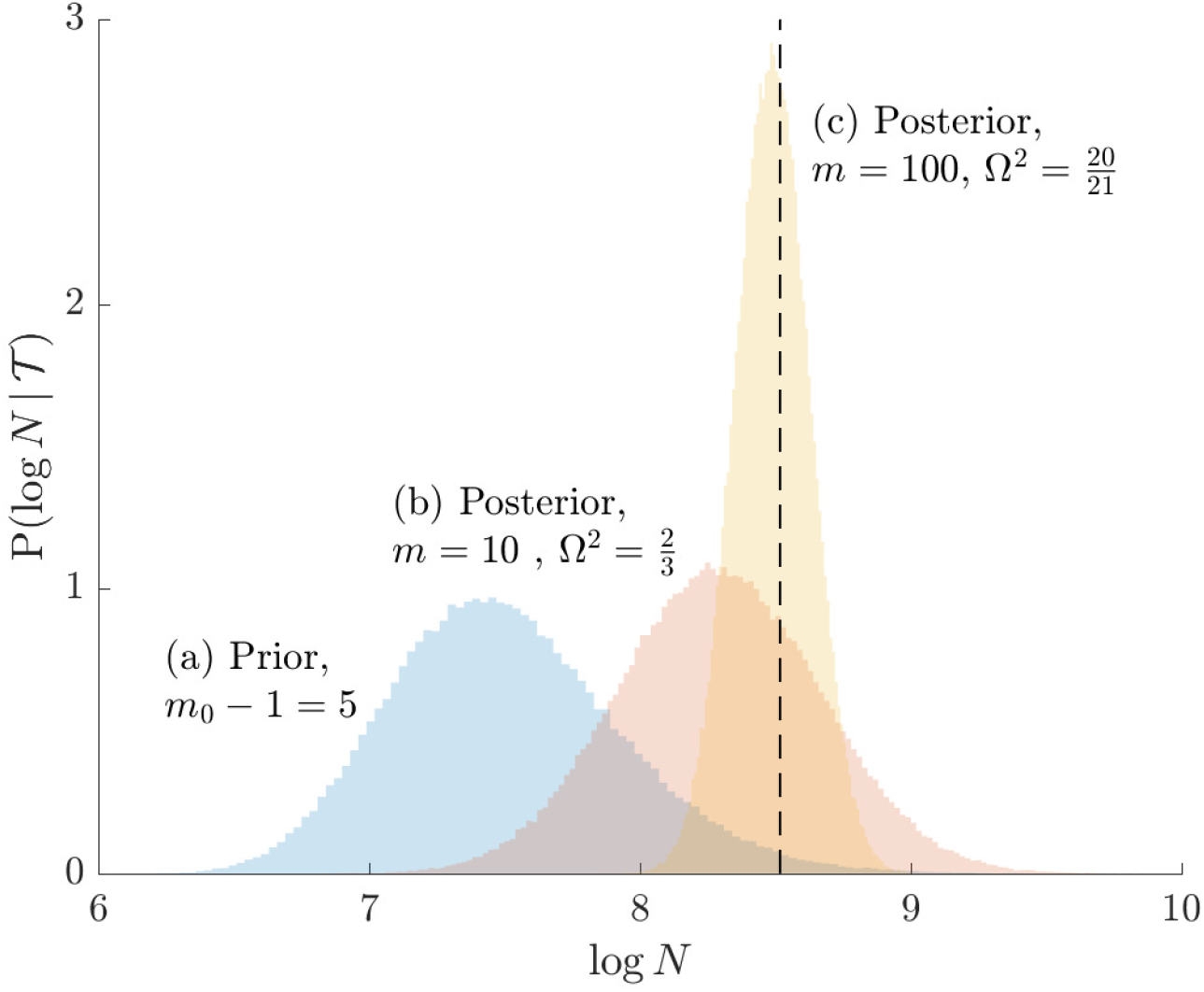
Effect of conjugate prior on Kingman coalescent estimation. We examine the relative impact on estimate precision of a conjugate Kingman prior that contributes *m*_0_ – 1 = 5 virtual observations. We work in log *N*_1_ for convenience. We compare this prior to posteriors, which are obtained under observed trees with *m* = 10 (red) and *m* = 100 (yellow) coalescent events. The true value is in black. The prior contribution decays as Ω^2^ increases towards 1.

### C. Skyline Smoothing Priors

In this section, we tailor Ω for the BSP, Skyride and Skygrid coalescent inference methods. These popular skyline-based approaches couple a piecewise-constant demographic coalescent data likelihood with a smoothing prior to produce population size estimates that change more continuously with time. The smoothing prior achieves this by assuming informative relationships between *N_j_* and its neighbouring parameters (*N*_*j*–1_, *N*_*j*+1_). Such *a priori* correlation implicitly introduces additional demographic information that is not available from the coalescent data 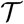. While these priors can embody sensible biological assumptions, we show that they may also engender overconfident statements or obscure parameter non-identifiability. We propose Ω as a simple but meaningful analytic for diagnosing these problems.

We first define uniquely objective (i.e. uninformative) reference skyline priors, which we denote P*(***ψ***). Finding objective priors for multivariate statistical models is generally non-trivial, but [3] state that if 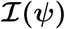 has form 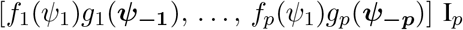 then 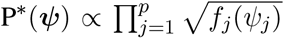. Here *f_j_* and *g_j_* are some functions and ***ψ_−j_*** symbolises the vector ***ψ*** excluding *ψ_j_*. Following this, we get

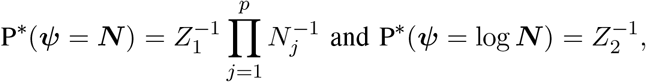

with *Z*_1_, *Z*_2_ as normalisation constants. Given its optimal properties [24], we only consider ***ψ*** = log ***N***, and drop explicit notational references to it. Under this parametrisation, 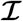 and its expectation with respect to the prior are equal, i.e. 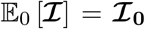. In addition, the reference prior in this case is 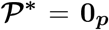, with **0_*p*_** as a matrix of zeros. This yields Ω = 1 by Eq. (4). A uniform prior over log-population space is hence uniquely objective for skyline inference.

Other prior distributions, which are subjective by this definition, necessarily introduce extra information and contribute to posterior estimate precision. This contribution will be reflected by an Ω < 1. The two most widely-used, subjective, skyline plot smoothing priors are:

i. the *Sequential Markov Prior* (SMP) used in the BSP [8], and
ii. the *Gaussian Markov Random Field* (GMRF) prior employed in both the Skyride and Skygrid methods [20] [11].

As the SMP and GMRF both propose nearest neighbour autocorrelations among elements of ***ψ***, tridiagonal posterior Fisher information matrices result. We represent these as 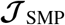 and 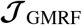, respectively.

The SMP is defined as: 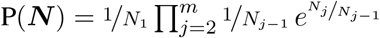 [8]. It assumes that 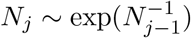 with a prior mean of *N*_*j*–1_. An objective prior is used for *N*_1_. To adapt this for log ***N***, we define 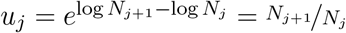 for *j* ∈ {1,…, *p* – 1}. In the Appendix we show how this expression yields Eq. (A1) and hence the transformed prior 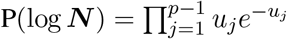. We then take relevant derivatives to obtain 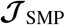, which for the minimally representative *p* = 3 case is written as:

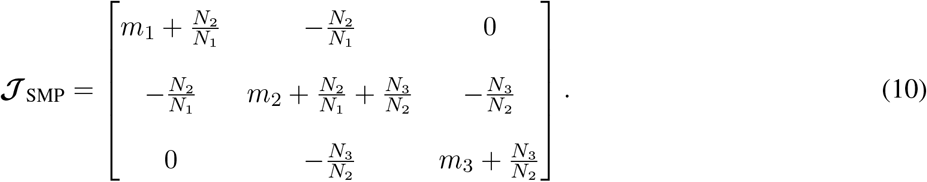

The *p* > 3 matrices simply extend the tridiagonal pattern of Eq. (10).

An issue with the SMP is its dependence on the unknown ‘true’ demographic parameter values. We cannot evaluate (or control) *a priori* how much information is contributed by this smoothing prior. Rapidly declining populations could feature *N*_*j*+1_/*N_j_* > *m_j_*, for example, which would result in prior over-reliance. Conversely, exponentially growing populations would be more data-dependent. This likely reflects the asymmetry in using sequential exponential distributions. The only control we have on smoothing implicitly emerges from choosing the number of segments, *p*. Some recent implementations of the BSP include an alternative log-normal prior that links *N_j_* with *N*_*j*–1_ [4], which is conceptually similar to the GMRF below.

The possibility of strong or inflexible prior assumptions under the BSP motivated the development of the GMRF for the Skyride and Skygrid methods [20]. The GMRF works directly with log ***N*** and models the autocorrelation between neighbouring segments with multivariate Gaussian distributions. The GMRF prior is defined as 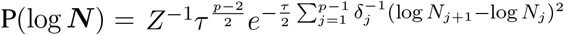 [20]. In this model, *Z* is a normalisation constant, *τ* a smoothing parameter, to which a gamma prior is often applied, and the *δ_j_* values adjust for the duration of the piecewise-constant skyline segments. Usually either (i) *δ_j_* is chosen based on the inter-coalescent midpoints in 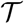 or (ii) a uniform GMRF is assumed with *δ_j_* = 1 for every *j* ∈ {1,…, *m* – 1}.

Similarly, we calculate 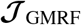 for the *p* = 3 case, which is:

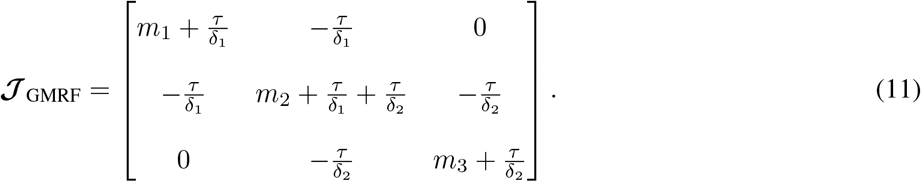

The Appendix provides the general derivation for any *p* ≥ 3. As *τ* is arbitrary and the *δ_j_* depend only on 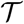, the GMRF is insensitive to the unknown parameter values. This property makes it more desirable than the SMP and gives us some control (via *τ*) of the level of smoothing introduced. Nevertheless, the next section demonstrates that this model still tends to over-smooth demographic estimates.

We diagonalise 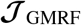 and 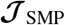 to obtain matrices of form 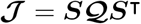. Here ***S*** is an orthogonal transformation matrix (i.e. |det [***S***]| = 1) and 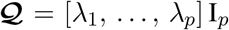 with λ*_j_* as the *j*^th^ eigenvalue of 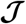. Since 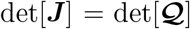, we can use Eq. (4) to find that 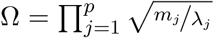. This equality reveals that λ*_j_* acts as a prior perturbed version of *m_j_*. When objective reference priors are used we recover *m_j_* = λ*_j_* and Ω = 1. We can use the ***S*** matrix to gain insight into how the GMRF and SMP encode population size correlations. The principal components of our posterior demographic estimates (which are obtained from 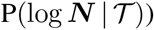 are the vectors forming the axes of the uncertainty ellipsoid described by 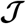.

These principal component vectors take the form {*e*_1_,…, *e_p_*} = {(log *N*_1_, 0,…, 0)^⊤^,… (0, 0,…, log *N_p_*)^⊤^} when we apply the reference prior P*(log ***N***). Thus, as we would expect, our uncertainty ellipses are centred on the parameters we wish to infer. However, if we use the GMRF prior these axes are instead transformed to {***S**e*_1_,…, ***S**e_p_*}. These new axes are linear combinations of log ***N*** and elucidate how smoothing priors share information (i.e. introduce autocorrelations) about log ***N*** across its elements. These geometrical changes also hint at how smoothing priors influence the statistical properties of our coalescent inference problem.

To solidify these ideas, we provide a visualisation of Ω and an example of ***S***. We consider the simple *p* = 2 case, where the posterior Fisher information and Ω for the GMRF and SMP both take the form:

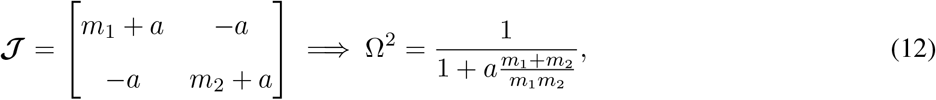

with *a* = *τ*/*δ*_1_ for the GMRF and *a* = *N*_2_/*N*_1_ for the SMP. The signal-to-noise ratio is 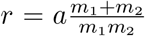 (see Eq. (9)) and performance clearly depends on how the *m* coalescent events in 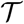 are apportioned between the two population size segments.

We can lower bound the contribution of these priors to Ω under any (*m*_1_, *m*_2_) settings by using the robust coalescent design from [24]. This stipulates that we define our skyline segments such that *m*_1_ = *m*_2_ = *m*/2 in order to optimise estimate precision under 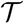. At this robust point we also find that max_{*m_j_*}_ Ω^2^ (or min_{*m_j_*}_ *r*) is attained. Fig. 2 gives the uncertainty ellipses for this robust *p* = 2 model at *a* = *m*/4. These are constructed in coordinates ***x*** = [*x*_1_,…, *x_p_*] centred about population size means log ***N*** as 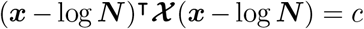 with *c* controlling the confidence level.

**Fig. 2:**
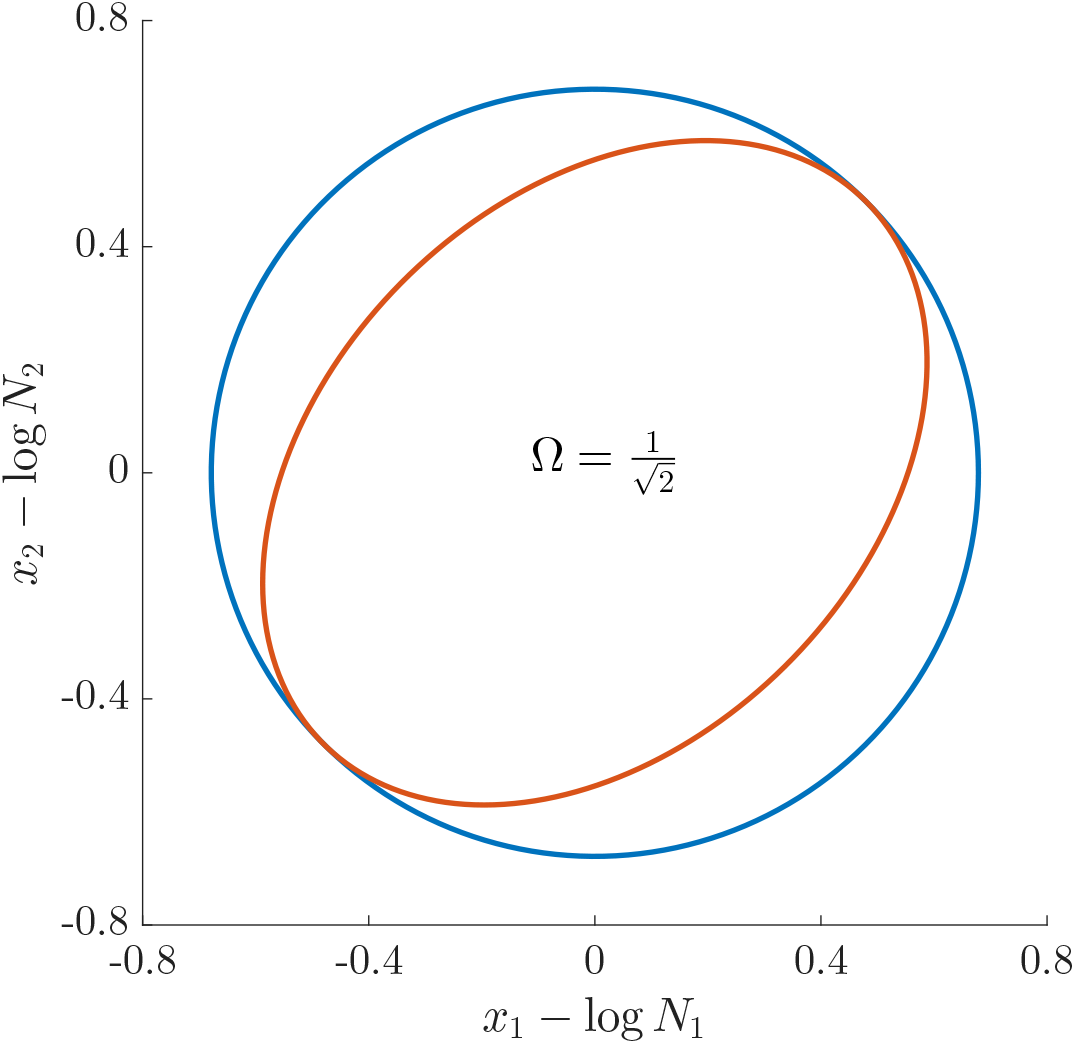
Uncertainty ellipses for SMP and GMRF. We show the improvement in asymptotic precision rendered by use of a smoothing prior for a *p* = 2 segment skyline inference problem. The prior informed ellipse (red) is smaller in volume and has skewed principal axes relative to the purely data informed one (blue). All ellipses represent 99% confidence with the *x_j_* indicating coordinate directions about their means, which are the log population sizes, log *N_j_*. The covariance that smoothing introduces controls the skew of these ellipses. Here Ω^2^ = 1/2, *m* = 40 (total coalescent event count) and *a* = 10 (this controls the prior influence see Eq. (12)). Larger *a* values lead to over-reliance on the smoothing prior.

Here 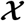 is either 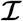 or 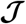. Because 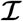 is diagonal the data-informed confidence ellipse has principal axes aligned with log ***N***. The covariance among population size segments in 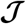, which is induced by the smoothing prior, skews these principal axes. We can see this by diagonalising 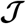 at *m*_1_ = *m*_2_ = *m*/2 and for every *r* to obtain:

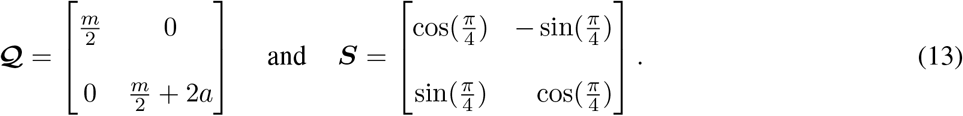

Applying ***S***, we find that the axes of our uncertainty ellipse (as visible in Fig. 2) have changed from 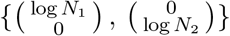 to 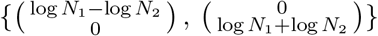 Sums and differences of log-populations are now the parameters that can be most naturally estimated under the SMP and GMRF. The reduction in the area of the ellipses of Fig. 2 is a proxy for Ω.

### D. The Dangers of Smoothing

Having defined ratios for measuring the contribution of smoothing priors to the precision of estimates, we now use them to explore and expose the conditions under which prior over-reliance is likely to occur in practice. We assume that skyline segments are chosen to satisfy the robust design *m_j_* = *m*/*p* for 1 ≤ *j* ≤ *p* [24], with *p* as the total number of skyline segments. We previously proved that robust designs, at *p* = 2, minimise dependence on the prior (maximise Ω). While this is not the case for *p* > 2, in Fig. A1 of the Appendix we illustrate that the maximal Ω point is generally well approximated by this robust setting. The Ω values computed here are therefore conservative for most {*m_j_*} settings. Other experimental designs rely more on the prior.

As in Eq. (5), we use the Ω^2^ = 1/2 threshold to diagnose when the coalescent data 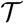 (likelihood) and prior are equally influencing demographic posterior estimate precision. At Ω^2^ = 1/2 the total Fisher information doubles since 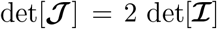. We previously uncovered the importance of this threshold in the Kingman conjugate prior problem, where it signified an equality between the number of pseudo and real samples contributed by the prior and data, respectively. As 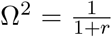 (see Eq. (8)), this setting is also meaningful because it achieves a unit signal-to-noise ratio for any skyline-based model.

We first reconsider the *p* = 2 case of Eq. (12), where *a* controls the prior contribution to 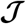. Here Ω^2^ = 1/2 suggests *a* = *m*/4, which implies that we are overly-reliant on smoothing when *a* is larger than 1/4 of the total observed coalescent events. This occurs when *N*_2_ ≥ *m*/4 *N*_1_ or *τ* ≥ *m*/4 *δ*_1_, for the SMP and GMRF respectively. The improved precision due to the prior at this *m*/4 threshold is shown in Fig. 2. The relative ellipse area (and hence Ω) will shrink further as we deviate from robust designs.

As the number of skyline segments, *p*, increase, smoothing becomes more influential and can promote misleading conclusions. For the *p* > 2 cases, we will only examine the GMRF, since the SMP has the undesirable property of dependence on the unknown *N_j_* values. To better expose the impact of the smoothing parameter *τ*, we will assume a uniform GMRF ({*δ_j_*} = 1) so that 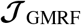 then only depends on {*m_j_*} and *τ*. We compute *r* and hence Ω, at various *p*. For example we find that

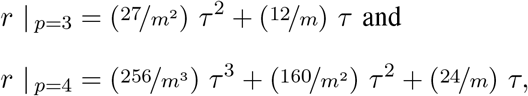

under the robust design. Interestingly, the order of the polynomial dependence of *r* (and hence Ω) on *τ* increases with *p*. We find that this trend holds for any {*m_j_*} design. We will use the term robust Ω for when Ω is calculated under a robust design.

Fig. 3 plots the robust Ω against *τ* and *p* for the uniform GMRF. A key feature of Fig. 3 is the steep *p*-dependent decay of Ω relative to the Ω^2^ = 1/2 threshold, which exposes how easily we can be unduly reliant on the prior, as *p* increases. Given a phylogeny 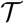, increasing the complexity of a skyline-based model enhances the dependence of our posterior estimate precision on the smoothing prior. This pattern is intuitive as fewer coalescent events now inform each demographic parameter [24]. However, Ω decays with surprising speed. For example, at *p* = 20 (the lowest curve in Fig. 3) we get Ω < 0.1 for *τ* = 1 and *m* = 100. Usually, *τ* has a gamma-prior with mean of 1 [20]. We show the corresponding mutual information increases due to these GMRF priors in Fig. A2 of the Appendix.

**Fig. 3:**
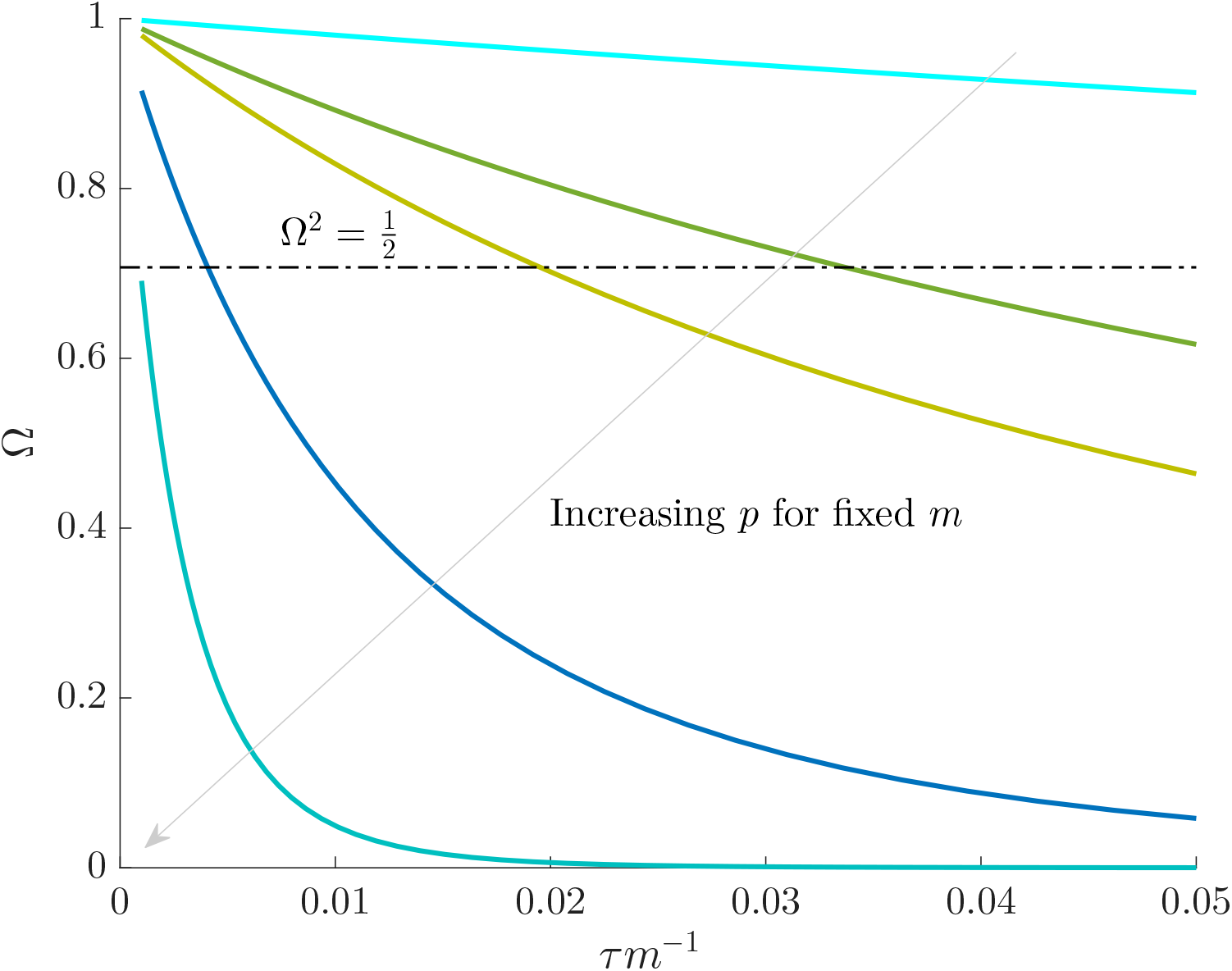
The impact of smoothing priors increases with skyline complexity. For the GMRF, we find that for a fixed *τ*/*m* (ratio of smoothing parameter to total coalescent event count), Ω significantly depends on the complexity, *p*, of our skyline. The coloured Ω curves are (along the arrow) for *p* = [2, 4, 5, 10, 20] at *m* = 100 with *m_j_* = *m*/*p* as the number of coalescent events per skyline segment. The dashed Ω^2^ = 1/2 line depicts the threshold below which the prior contributes more than the coalescent data to posterior estimate precision (asymptotically). For a given tree and *τ*, the larger the number of demographic parameters we choose to estimate, the stronger the influence of the prior on those estimates.

While Fig. 3 might seem specific to the uniform GMRF, it is broadly applicable to the BSP, Skyride and Skygrid methods. We now outline the implications of Fig. 3 for each of these skyline-based approaches.

#### a) (1) Bayesian Skyline Plot

This method uses the SMP, which depends on the unknown *N_j_* values. However, the results of Fig. 3 are valid if we set *τ* to 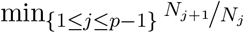, which results in the smallest non-data contribution to Eq. (10). This follows as 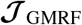 and 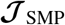 have similar forms. While this choice underestimates the impact of the SMP, it still cautions against high-*p* skylines and confirms suspected BSP issues related to poor estimation precision when skylines are too complex, or the coalescent data are not sufficiently informative [13]. However, good use of the BSP grouping parameter [8], which sets *p* < *m*, could alleviate these problems.

#### b) (2) Skyride

When this method uses the uniform GMRF, all results apply exactly. In its full implementation, the Skyride employs a time-aware GMRF that sets *δ_j_* based on 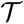 and estimates *τ* from the data [20]. However, even with these adjustments, the GMRF can over-smooth, and fail to recover population size changes [13, 9]. Our results provide a theoretical grounding for this observation. The Skyride constrains *p* = *m* and then smooths this noisy piecewise model. Consequently, it constructs a skyline which is too complex by our measures (the lowest curve in Fig. 3 is at *p* = *m*/5). By rescaling the smoothing parameter to 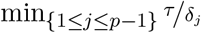, the Ω curves in Fig. 3 upper bound the true Ω values of the time-aware GMRF.

#### c) (3) Skygrid

This method uses a scaled GMRF. For a tree with TMRCA *T*, the Skygrid assumes new population size segments every *T*/*p* time units [11]. As a result, every *δ_j_* = *T*/*p* and the time-aware GMRF becomes uniform with rescaled smoothing parameter *τ*/*p*. Therefore, the conclusions of Fig. 3 hold exactly for the Skygrid, provided the horizontal axis is scaled by *p*. This setup reduces the rate of decay but the Ω curves still caution strongly against using skylines with *p* ≈ *m*. Unfortunately, as its default formulation sets *p* to 1 less than the number of sampled taxa (or lineages) [11], the Skygrid is also be vulnerable to prior over-reliance.

The popular skyline-based coalescent inference methods therefore all tend to over-smooth, resulting in population size estimates that can be overconfident or misleading. This issue can be even more severe than Fig. 3 suggests since in current practice *p* is often close to *m* and non-robust designs are generally employed. Further, skylines are only statistically identifiable if every segment has at least 1 coalescent event [24, 22]. Consequently, if *p* > *m* is set, smoothing priors can even mask identifiability problems. We recommend that 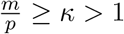 must be guaranteed and in the next section derive a model rejection guideline for finding κ, the suggested minimum number of coalescent events per skyline segment, and diagnosing prior over-reliance.

### E. Prior Informed Model Rejection

We previously demonstrated how commonly-used smoothing priors can dominate the posterior estimate precision when coalescent inference involves complex, highly parametrised (large-*p*) skyline models. Since data are more influential than the prior when Ω^2^ > 1/2, we can use this threshold to define a simple *p*-rejection policy to guard against prior over-reliance. Assume that the 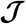 matrix resulting from our prior of interest is symmetric and positive definite. This holds for the GMRF and SMP. The standard arithmetic-geometric mean inequality, 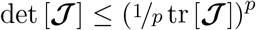, then applies with tr denoting the matrix trace. Since 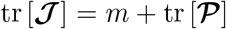 we can expand this inequality and substitute in Eq. (4) to get 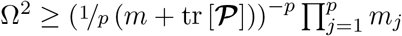.

Since this inequality applies to all {*m_j_*}, we can maximise its right hand side to get a tighter lower bound on Ω^2^. This bound, termed *ω*^2^, is achieved at the robust design *m_j_* = *m*/*p* and is given by

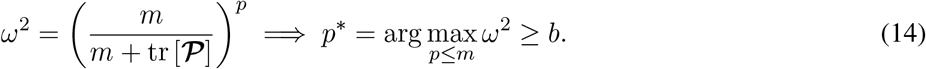

We define *b* ≥ 1/2 as a conservative model rejection criterion with *ω*^2^ ≥ *b* implying that Ω^2^ ≥ *b*. If *p*\* is the largest *p* satisfying these inequalities (see Eq. (14), arg indicates argument), then any skyline with more than *p*\* segments is likely to be overly-dependent on the prior and should be rejected under the current coalescent tree data.

Alternatively, we recommend that skylines using a smoothing prior (with matrix 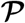) should have at least *κ* = *m*/*p*\* events per segment to avoid prior reliance. The *p* ≤ *m* condition in Eq. (14) ensures skyline identifiability [24] and generally *p*\* ≤ *m*/2 (i.e. *κ* > 1). The dependence of *ω*^2^ on 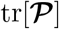 means that additions to the diagonals of 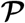 necessarily increase the precision contribution from the prior. This insight supports our previous analysis, which used *τ* from the uniform GMRF to bound the performance of the SMP and time-aware GMRF. In the Appendix (see Eq. (A2)) we derive analogous rejection bounds based on the excess mutual information, 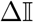, from Eq. (7). There we find that *p* acts like an information-theoretic bandwidth, controlling the prior-contributed mutual information.

Eq. (14), which forms a key contribution of this work, can be computed and is valid for any smoothing prior of interest. For the uniform GMRF where 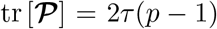, we get 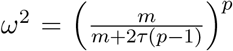. Note that *ω*^2^ = 1 here whenever *p* = 1 or *τ* = 0, as expected (i.e there is no smoothing at these values). In Fig. A4 of the Appendix, we confirm that *ω*^2^ is a good lower bound of Ω^2^. We enumerate *ω*^2^ across *τ* and *p*, for an observed tree with *m* = 100, to get Fig. 4, which recommends using no more than *p*\* = 19 segments (κ ≈ 5.3). In Fig. A5 we plot *p*\* curves for various *m* and *τ*, defining boundaries beyond which skyline estimates will be overly-dependent on the GMRF.

**Fig. 4:**
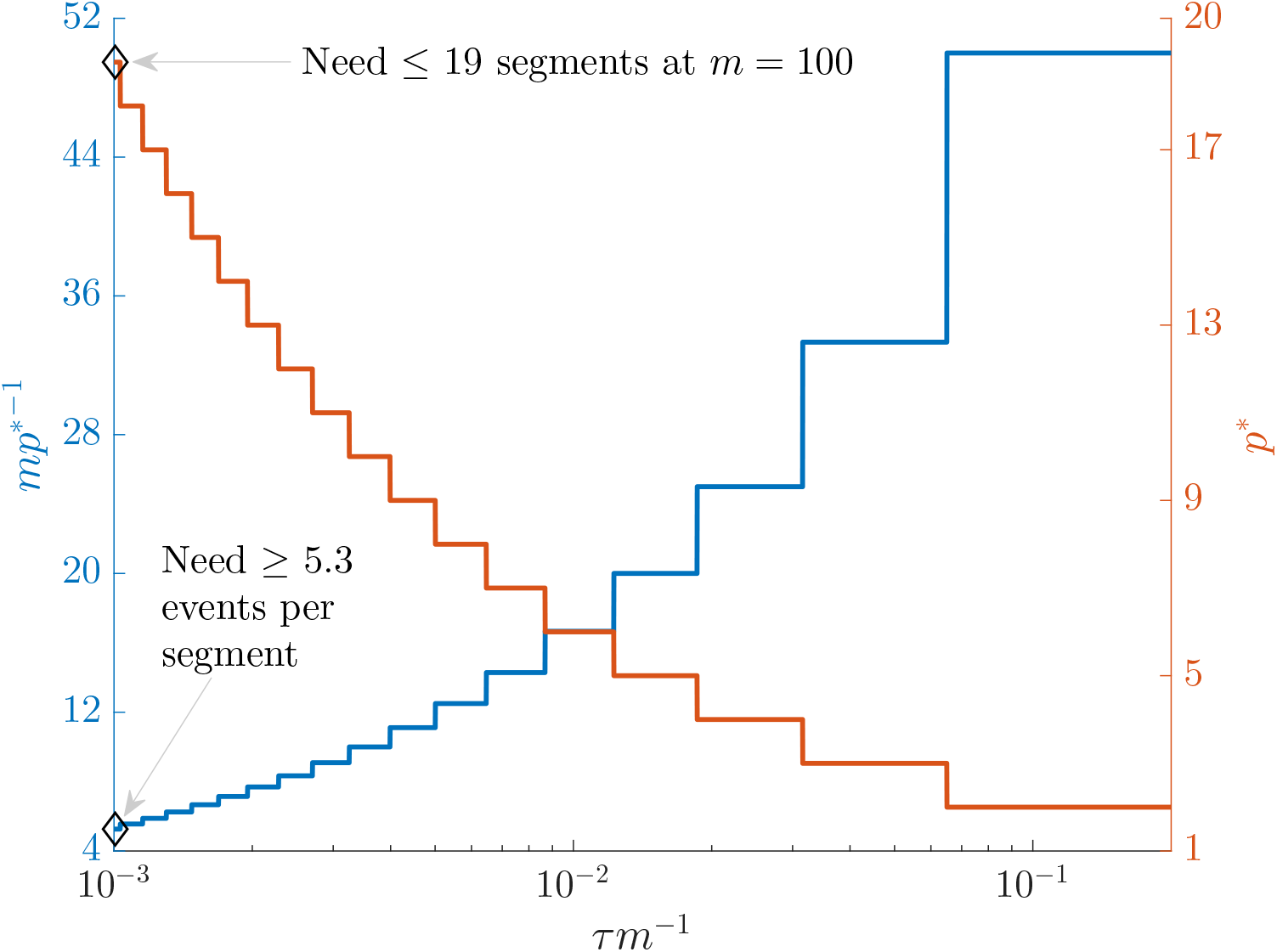
Bounding skyline complexity using the prior-data tradeoff. For the GMRF with uniform smoothing, we show how the maximum number of recommended skyline segments, *p*\* (red), decreases with prior contribution (level of smoothing i.e. increasing *τ*/*m*). Hence the minimum recommended number of coalescent events per segment, *κ* = *m*/*p*\* (blue), rises. Here we use the *ω*^2^ ≥ *b* = 1/2 boundary (Eq. (14)), which approximates Ω^2^ and provides a more easily computed measure of prior-data contributions. At larger *b* the *p*\* at a given *τ*/*m* decreases. The *p*\* measure provides a model rejection tool, suggesting that models with *p* > *p*\* should not be used, as they would risk being overly informed by the prior.

In the Appendix we further analyse Eq. (14) for the uniform GMRF to discover that Ω^2^ is bounded by curves with exponents linear in *τ* and quadratic in *p* (see Eq. (A3)). This explains how the influence of smoothing increases with skyline complexity and yields a simple transformation 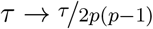, which can negate prior over-reliance. For comparison, the *Skyride* implements *τ* → *τ*/*p*. The marked improvement, relative to Fig. 3, is striking in Fig. A3. Other revealing prior-specific insights can be obtained from Eq. (14), reaffirming its importance as a model rejection statistic.

Our model rejection tool of Eq. (14) can serve as a useful diagnostic for skyline over-parametrisation, and as a precaution against prior over-reliance. However, we do not propose *p*\* as the sole measure of optimal skyline complexity; because while *p*\* warns against the prior being too relatively influential, it does not guarantee any absolute estimate precision e.g. a small (*m*, *τ*) pair might produce the same *p*\* as a larger pair. Choosing an optimal *p* in a data-justified manner is an open problem that is still under active study [21]. We next illustrate how Ω^2^, via its more easily computed approximation, *ω*^2^, can be practically applied to detect and reject over-smoothed skyline plot models, using datasets that are commonly employed to evaluate the performance of coalescent demographic inference.

### F. Illustrative Examples: Egyptian HCV and Beringian Bison

We validate the practical utility of *ω*^2^ (and hence Ω^2^), as a diagnostic of prior over-dependence, by investigating changes in effective population size inferred from the well-studied Egyptian HCV-4 [25] and Beringian steppe bison [31] datasets. The first consists of 63 partial sequences of HCV genotype 4 and was previously analysed in [25] using a coalescent model with a parametric demographic function that featured periods of constant population size separated by a phase of exponential growth. The second dataset comprises 152 modern and partial mtDNA and was investigated in [31], where skyline plot models confirmed a demographic history of exponential growth then decline (known as a boom-bust trend). These two datasets have since been re-examined under various alternate models in [20, 11, 22] and several other studies.

We simulated 100 trees with *m* + 1 = *n* = 63 and 152 tips, using the software package MASTER [37], according to inferred HCV and bison population size trends respectively. The HCV population size trend that we simulated from is provided in [25]. We inferred the population size trend of the bison dataset using the BSP (with sequential Markovian prior) in accordance with published analyses [8]. We used 20 population groups and the optimal design from [24] to ensure that we captured complex bison population dynamics reliably. As our focus is on exploring the behaviour of skylines and *ω*^2^ given a particular underlying population size trend and not the uncertainty associated with that trend, we used the posterior mean (HCV) or median (bison) of these inferred trends for simulating trees and do not consider genealogical uncertainty.

The simulated set of coalescent trees from each dataset provide an approximate measure of the coalescent variance that could arise from the inferred underlying population size trends. We then estimated log ***N*** from every simulated tree using various skyline models with time-aware GMRF smoothing priors, as in [20]. We varied the relative contributions of the coalescent data and GMRF to our posterior log-population size estimates by changing either the skyline dimension, *p*, or the GMRF smoothing parameter *τ*. As *m* is fixed for a given dataset and robust designs are applied, increasing the number of coalescent events in each segment, *m_j_*, reduces *p*.

We analysed every tree over all combinations of *m_j_* ∈ {1, 2, 4, 8} across a wide range of *τ*. For comparison, we also generated purely data-informed estimates of log ***N***, for the same *m_j_*, by replacing the subjective GMRF with a uniform, objective prior. We computed *ω*^2^ from Eq. (14) for these settings in Fig. 5 and observe that, as expected, it decreases with both *τ* and *p* (i.e. *ω*^2^ increases with *m_j_*). Practical analyses of these datasets using Skyride or Skygrid approaches, would choose or infer a *τ* value and set *p* ≈ *m*. However, Fig. 5 shows *κ* = *m*/*p*\* > 1 and hence *m_j_* > 1 events per skyline parameter are often necessary to achieve *ω*^2^ ≥ 1/2. This raises questions about the validity of the common practice of applying these methods using their default settings.

**Fig. 5:**
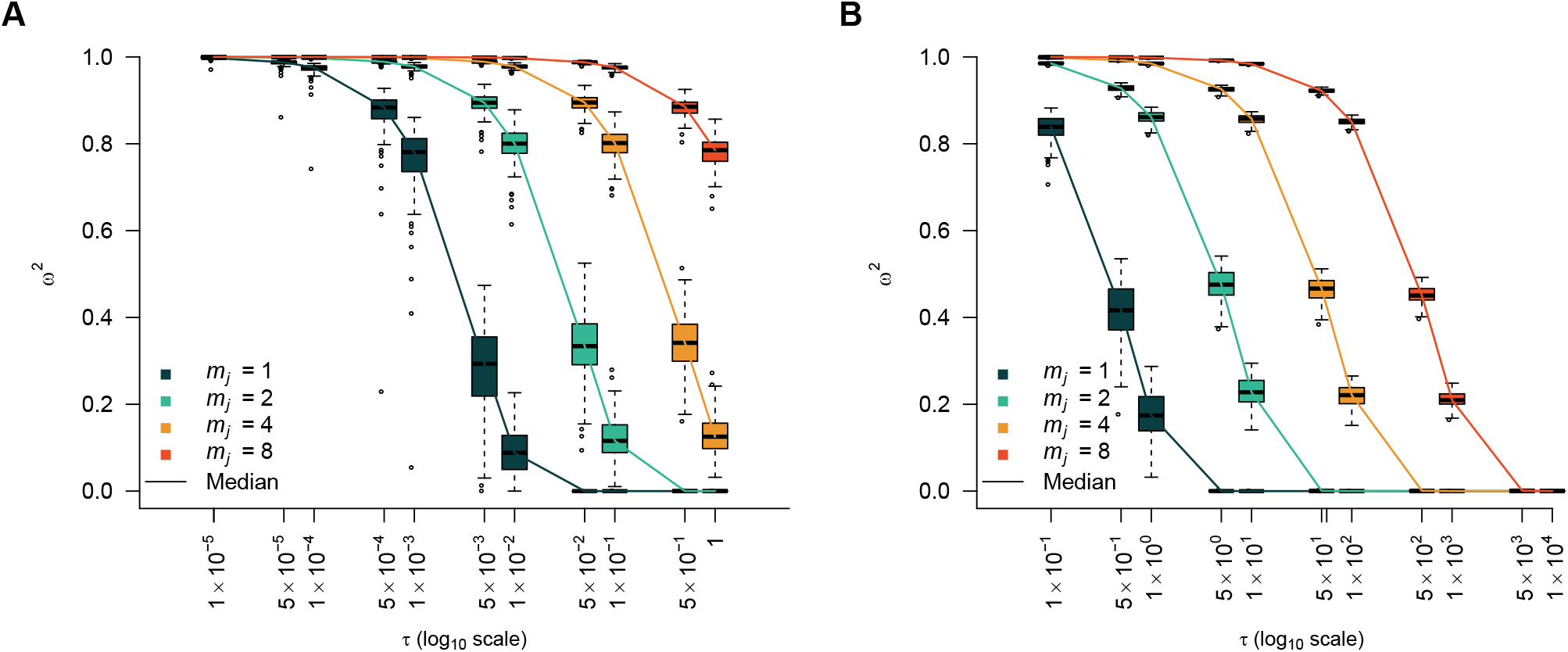
Model rejection statistics for the HCV and bison datasets. The metric *ω*^2^ is calculated for each tree (see Eq. (14)) under a time-aware GMRF for various combinations of its smoothing parameter *τ* and *m_j_*, the number of coalescent events per skyline segment. The box-plots summarise the resulting *ω*^2^ over 100 simulated trees that represent the demographic histories of the (A) Egyptian HCV and (B) Beringian bison datasets. The solid lines link the median values across boxes for a given *m_j_* and hence skyline dimension *p* (*m_j_* = *m*/*p*). We discourage the use of skyline models with *ω*^2^ < 1/2.

Fig. 5 confirms that the recommended maximum skyline dimension *p*\* falls and hence the minimum allowable number of coalescent events per segment *m_j_* grows as the smoothing parameter *τ* increases. We demonstrate the qualitative difference in skyline-based estimates between *p* values on either side of the *p*\* criterion for a single simulated HCV and bison tree in Fig. 6. In panels A and C we present the Skyride estimate, which uses *m_j_* = 1 and implements *p* > *p*\*, at the chosen *τ* values (0.05 and 1). Contrastingly, in B and D, we illustrate an equivalent skyline with a different *m_j_*, which achieves *p* < *p*\* at this same *τ*, according to our *ω*^2^ metric (see the *m_j_* = 4 and *m_j_* = 2 curves at *τ* = 0.05 and 1 in panels A and B of Fig. 5) respectively). We overlay the corresponding skyline (with the same *m_j_*) obtained with an objective uniform prior, to visualise the uncertainty engendered from the coalescent data alone.

**Fig. 6:**
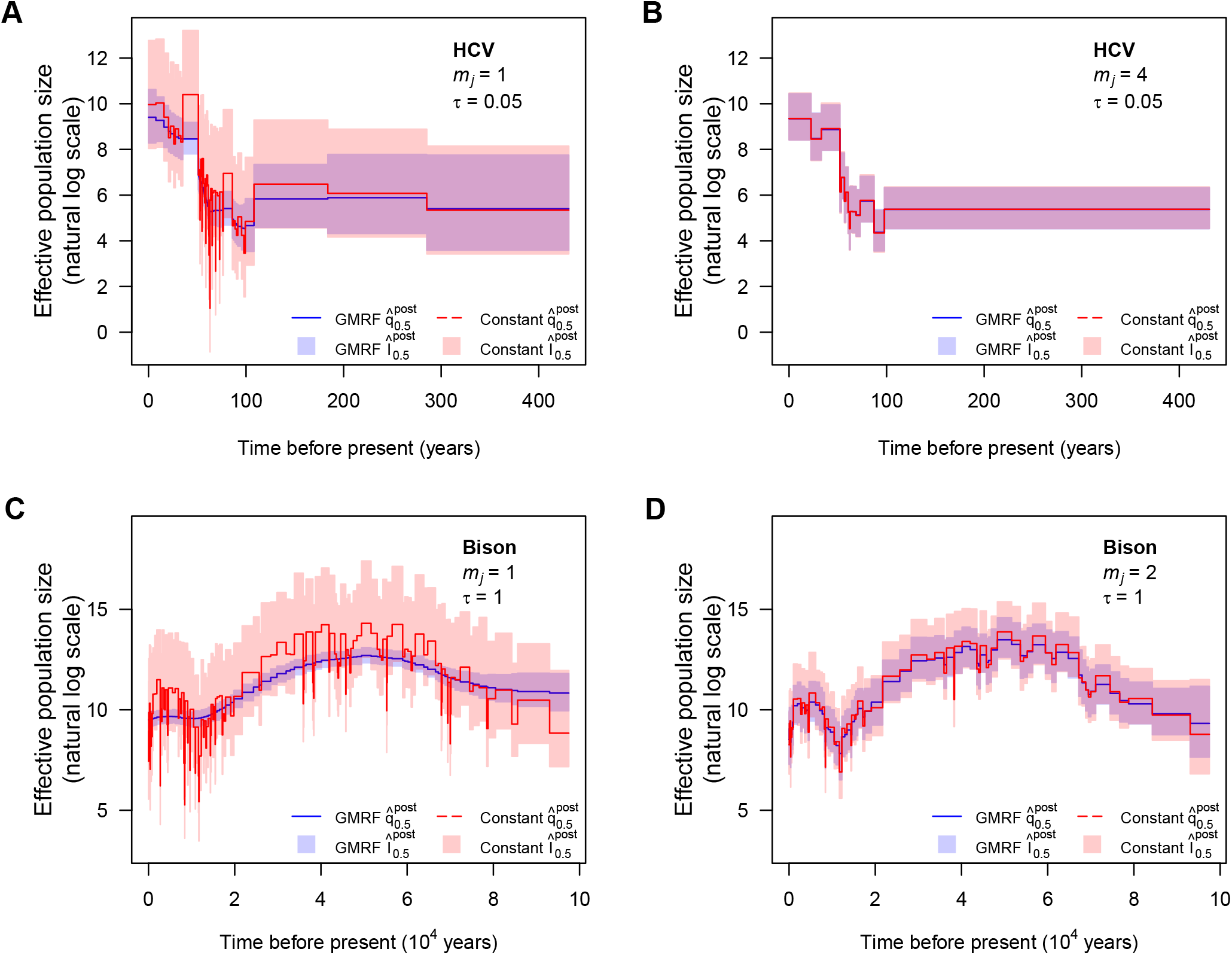
HCV and bison demographic estimates under GMRF and uniform priors. We analyse demographic estimates under time-aware GMRF priors (blue) and objective uniform priors (red) for a single tree simulated under the demographic scenarios inferred from the Egyptian HCV (A and B) and Beringian bison (C and D) datasets. In panels A and C we present Skyride estimates, which use *m_j_* = 1 and *τ* = 0.05 (A) and 1 (C). These skylines have dimension *p* that is larger than our maximum recommended dimension *p*\*, which is computed from Fig. 5. In panels B and D we re-estimate population size at *m_j_* = 4 (B) and 2 (D). These groupings of coalescent events achieve *p* < *p*\* as justified by our *ω*^2^ metric (see Eq. (14)). Solid lines are posterior medians while semi-transparent blocks are the 95% HPD intervals.

At *m_j_* = 1 (panels A and C of Fig. 6), the uniform prior produces a skyline that infers more rapid demographic fluctuations through time than that estimated with the GMRF prior. Further, the 95% HPD intervals from the uniform prior (red) are substantially wider than those from the GMRF prior (blue) in both examples, highlighting the marked contribution of the time-aware GMRF prior to posterior estimate precision. While this smoothed trajectory looks reliable we argue that, because *p* > *p*\* (and hence 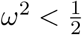), it is difficult to justify using the data alone and that the prior is responsible for too much of the estimate precision. In contrast, at *m_j_* = 4 and *m_j_* = 2 (panels B and D of Fig. 6), which apply *p* < *p*\*, both prior distributions yield more similar skylines, implying that GMRF smoothing has not substantially inflated posterior estimate precision.

Under these settings we have fewer demographic fluctuations than for *m_j_* = 1 because 4 and 2 times more coalescent events are informing each parameter or skyline segment, respectively. We achieve smaller uncertainty than *m_j_* = 1 with a uniform prior (which is overfitted) but without excessively relying on the GMRF smoothing, which at *m_j_* = 1 is likely underfitting. The *ω*^2^ metric and hence *p*\* criterion help us better balance data, noise and our prior assumptions. In contextualising these results it is important to note that skyline plots provide harmonic mean and not point estimates of population size [26]. Consequently, we are inferring sequences of means from our coalescent data, which *a priori* may not need to conform to a smooth pattern.

The HCV example shows that for times beyond *t* > 100 years there are so few events that it is more sensible to estimate a single mean (panel B), which we are confident in across this period, as opposed to several less certain and overfitted means (panel A). In contrast, for the bison example, the bottleneck over 10^4^ < *t* < 2 × 10^4^ years is oversmoothed (panel C), despite many coalescent events occurring in that region. The simple correction of extending our harmonic mean over 2 events (panel D) restores the necessary fall in population size. Deciding on how to balance uncertainty with model complexity is non-trivial and, as shown in these examples, caution is needed to avoid misleading conclusions. We posit that *ω* (and hence Ω) can help formalise this decision-making and improve our quantification of the uncertainty across skyline plots.

Having confirmed Ω as a credible measure of relative uncertainty, we briefly explore how it relates to more easily ascertained measures of uncertainty. For each simulated coalescent tree in the HCV example above we computed Ω (via Eq. (4)) and two ancillary statistics based on the 95% highest posterior density (HPD) intervals of the log ***N*** estimates. These are the median HPD ratio q_0.5_ and the relative HPD product (across the skyline segments) 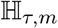, which are formulated as:

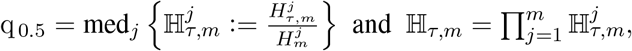

with med indicating the median value of a set. Here 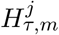 is the 95% HPD interval of log *N_j_* under a GMRF with smoothing parameter *τ* and 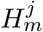 is the equivalent HPD when the objective uniform prior is applied instead.

The 95% HPD interval is closely connected to the inverse of the Fisher information matrices that define Ω and, further, describes the most visually conspicuous representation of the uncertainty present in skyline plot estimates. Comparing Ω to these ancillary statistics, which evaluate the median and total 95% uncertainty of a skyline plot, allows us to contextualise Ω against more relatable (though different) and obvious visualisations of posterior performance. We present these comparisons in Fig. A6 of the Appendix. There we find that all statistics monotonically decay with *τ* i.e. as the time-aware GMRF becomes more informative. The sharpness of this decay is highly sensitive to *m_j_*. Larger *m_j_* means that more coalescent data are informing each estimated parameter (smaller *p*).

The reduced decay with *m_j_* supports our assertion that *p* acts as an exponent controlling prior over-reliance (see Fig. 3). The gentler decay of q_0.5_ (relative to Ω and 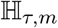), which largely does not account for *p*, confirms that we could be misled in our understanding of the impact of smoothing if we neglected skyline dimension. In contrast Ω and 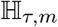, which both measure, in some sense, the relative volumes of uncertainty across the entire skyline-plot due to the data alone and the data and prior, fall more significantly and consistently. At *m_j_* = 1 (*p* = *m*), which is the most common setting in the Skyride and Skygrid methods, both statistics are markedly below 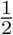 and posterior estimates will often be too dependent on the prior. This high-*p* behaviour is also indicative of model over-parametrisation [21]. Our metric Ω therefore relates sensibly to visible and common proxies of uncertainty.

## III. Discussion

Popular approaches to coalescent inference, such as the BSP, Skyride and Skygrid methods, all rely on combining a piecewise-constant population size likelihood function with prior assumptions that enforce continuity. This combination, which is meant to maximise descriptive flexibility without sacrificing the smoothness that is expected to be exhibited by real population size curves over time, has led to many insights in phylodynamics [13]. However, it has also spawned concerns related to over-smoothing and lack of methodological transparency [20] [9]. In this work we attempted to address these concerns by deriving metrics for diagnosing and clarifying the existing assumptions present in current best practice.

Detecting and correcting for underfitting or over-smoothing is crucial if reliable and meaningful assessments of the effective population size changes of a species or pathogen of interest are to be made from sequence data. Abrupt changes in effective population size are not only biological plausible but may also signal key events that have shaped the demographic histories of populations [27]. In ecology, identifying mass extinctions and bottlenecks in diversity might signify the impact of rapid environmental change or anthropogenic influences (e.g., hunting or changes in land use). Similarly, in epidemiology, sharp fluctuations in the prevalence of an infection might support hypotheses about emergence in novel populations, seasonality, the effect of interventions, vaccines, or drug treatments. Further, rapid exponential growth of any population may, when observed over a longer timescale, appear as a near-stepwise transition in population size.

Underfitting these changes would limit understanding of the dynamics of the study population and could affect conclusions about the potential causative factors that influenced those dynamics. However, recognising when commonly used methods for inferring these demographic trends are over-smoothing is difficult. By capitalising on (mutual) information theory and (Fisher) information geometry we formulated the novel coalescent information ratio, Ω, which provides a rigorous means of solving this overs-moothing problem. This ratio describes both the proportion of the asymptotic uncertainty around our posterior estimates that is due solely to the data and the additional mutual information that the prior assumptions introduce.

We derived analytic expressions for Ω for the BSP, Skyride and Skygrid estimators of effective population size, which combine piecewise skyline likelihoods with either SMP or GMRF smoothing priors. We also showed that Ω has an exact and intuitive interpretation as the ratio of real coalescent events to the sum of real and virtual (prior-contributed) ones in a Kingman coalescent model. Using Ω^2^ = 1/2 as a threshold delimiting when the prior contributes as much information as the coalescent data, we found that it is easy to become overly dependent on prior assumptions as the skyline dimension, *p*, increases (for a fixed tree size). This central result emerges from the drastic reduction in the number of coalescent events informing on any population size parameter as *p* rises. Per parameter, the BSP and Skyride use only a few or one event respectively [20, 8], while the Skygrid may have no events informing some parameters [11].

These issues can be obscured by current Bayesian implementations, which can still produce apparently reasonable population size estimates, at least visually, as illustrated in our simulated HCV and bison case studies. Our simulations indicate that analyses that combine maximally parametrised skylines (one event per segment or parameter) with GMRF smoothing can lead to errors in population size inference. For trees simulated according to the HCV demographic scenario, estimates were likely overfitted in the far past, inflating HPDs, but oversmoothed towards the present. The resulting skyline uncertainty contrasted that from the original [25] and later [23] analyses. In the bison example, we found evidence for underfitting. The inferred skyline there emphasised a smoother boom-bust trend with concentrated HPDs. However, this underestimated the depth of a bottleneck during which coalescent events were concentrated.

These mismatches between data and smoothing can be difficult to diagnose and problematic, not just for prior over-dependence. Low coalescent event counts, for example, can lead to poor statistical identifiability [30] which might manifest in spurious MCMC mixing. Consequently, we proposed a practical *p*\* rejection criterion for ensuring that coalescent data is the main source of inferential information. This criterion, which was based on an approximation to Ω^2^, provided a way of regularising skyline complexity. When applied to our examples it recommended a 4-event skyline grouping that resulted in demographic reconstructions that were more consistent with the above mentioned HCV studies. It also suggested a simple 2-event grouping that recovered the bison bottleneck dynamic without generating too much estimate noise.

This *p*\* criterion bounds the maximum recommended skyline dimension for a given dataset (tree) size and provides a usable means of defining the minimum number of coalescent events, *κ*, which we should allocate to each skyline segment to guard against too much prior influence. Since *κ* only requires our computing the sum of the diagonals of the prior Fisher matrix, it can serve as a simple rule-of-thumb for sensibly balancing the prior-data tradeoff in skyline plots (e.g. in the BSP, the grouping parameter might be set to a value above *κ* to ensure well-regularised estimates). As we found Ω^2^ to be lower-bounded by more visible measures of skyline uncertainty, such as the product of relative HPD widths, useful approximations to *p*\* and *κ* may also be computed from these measures.

Our Ω metric also provides insight into how we can alleviate the dramatic impact of skyline complexity on prior over-reliance. When specialised to the GMRF, for example, it reveals that we can negate over-smoothing by scaling the smoothing parameter *τ* with a quadratic of *p*. Moreover, it shows that only by increasing the information available from the sampled phylogeny can we reasonably allow for more complex piecewise-constant functions under a given prior. Recent methods, such as the *epoch sampling skyline plot* [22], which can double the Fisher information extracted from a given phylogeny by exploiting the informativeness of sampling times, would support higher dimensional skylines. Such approaches have the potential to increase the contribution of the data without elevating the influence of the smoothing prior.

While in this paper we have applied Ω to non-parametric, skyline inference problems in population genetics, ecology and epidemiology, its general formulation in Eq. (4) is more widely applicable. It can be also applied to coalescent inference problems where specific parametric models (e.g., exponential/logistic growth) are used, in order to disentangle the contributions of observed data and the prior distributions over these parameters, though numerical solutions will likely be necessary. More generally, our approach is valid for any statistical problem, provided the Hessian matrices necessary for deriving the prior and data Fisher information terms are valid and computable. This is not limited to prior-data tradeoffs. Similar ratio metrics should be derivable by comparing Fisher information terms from different sources (e.g. to test whether one source of data is more informative than another).

Thus, we have devised and validated a rigorous means of better understanding, diagnosing and preventing prior over-dependence. We hope that our statistic, which clarifies and quantifies the often inscrutable impact of the prior and data, will help researchers make more active and considered design decisions when adapting popular skyline-based techniques. Our work also aligns with recent studies, which have started to re-examine both model selection and prior definition [21, 9] in an attempt to derive more reliable effective population size estimates from coalescent trees. While we believe that data-driven conclusions are generally the most justifiable we note that, in the context of skyline plots, this can be open to interpretation and the choice of prior is far from trivial.

## Acknowledgments

We thank Louis du Plessis for his useful comments and insights on this project. This study was funded by the UK Medical Research Council (MRC) and the UK Department for International Development (DFID) under the MRC/DFID Concordat agreement and is also part of the EDCTP2 programme supported by the European Union (grant reference MR/R015600/1). This work was supported by the Oxford Martin School.

## Supplementary Material

Data (and code in Matlab) available from the Dryad Digital Repository: https://datadryad.org/stash/dataset/doi.10.5061/dryad.1jwstqjs2

## Appendix

### A. Smoothing Prior Fisher Information Matrices

Here we derive the prior-informed Fisher information matrices for the SMP and GMRF smoothing priors. We start by finding the log-population size transformed version of the SMP smoothing prior. We then calculate its Hessian to get 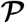, and so obtain the general form of Eq. (10). The SMP is given in [8] as 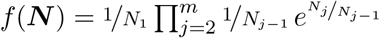. We define ***η*** = *ρ*(***N***) := log ***N*** so that its inverse *ρ*^−1^(***η***) = *e^**η**^*. These expressions are in vector form so ***η*** = [*η*_1_,…, *η_p_*] = [log *N*_1_,…, log *N_p_*]. We want the transformed prior *g*(***η***). Applying the multivariate change of variables formula gives *g*(***η***) = *f*(*e^**η**^*)|det [Δ*ρ*^−1^]|, with Δ*ρ*^−1^ = [*e*^*η*_1_^,…, *e^η_p_^*] **I**_*p*_ as the Jacobian of *ρ*^−1^. This implies that 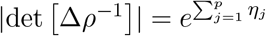. Substituting and expanding gives the SMP log-prior:

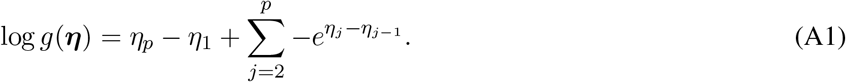

We can then obtain 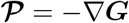, with ***G*** = log *g*(***η***). The diagonals of 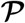 are: 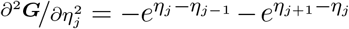 for 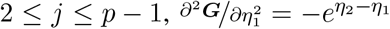 and 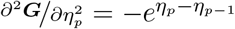. The non-zero off-diagonal terms are: 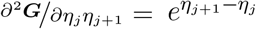 and 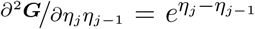. The result is a symmetric tridiagonal matrix that has zero row and column sums. The 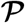 matrix is then added to the Fisher information matrix 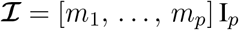 (with *m_j_* as the number of coalescent events informing on the *j*^th^ parameter), to get 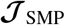.

We now compute 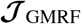 which is given in the main text as Eq. (11). For the GMRF 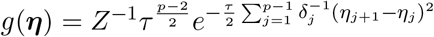 [20] and so 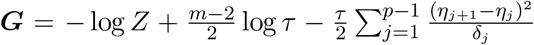. Taking second derivatives we get diagonal terms of the Hessian, **∇*G***, as: 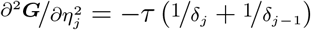 for 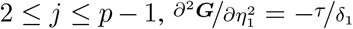 and 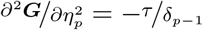. The non-zero off diagonal terms are: 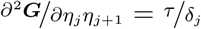 and 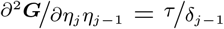. The GMRF also gives a symmetric tridiagonal 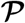 with row and column sums of zero. Adding −**∇*G*** to the diagonal 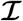 matrix yields 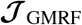.

### B. Further Smoothing Results

In the main text we asserted that the Ω computed at the robust point of *m_j_* = *m*/*p* [24] generally upper bounds the achievable Ω values at other *m_j_* settings. Here we provide evidence for this assertion. While strictly arg max_{*m_j_*}_ Ω ≠ *m*/*p* (except for *p* = 2), we numerically find that 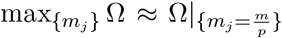. We show this for the GMRF under uniform smoothing in Fig. A1. This makes sense as while (for fixed smoothing parameters) arg 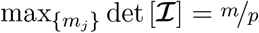 and 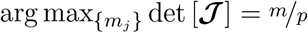, there is no reason to believe that this also maximises their ratio. The sawtooth Ω curves in Fig. A1 reflect changes in the other {*m_j_*} values, given a fixed *m*_1_.

Hence we used the robust design point in our calculation of the Ω^2^ curves for the GMRF in Fig. 3. The corresponding additional mutual information 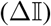 curves for this case are provided in Fig. A2. These show how larger values of the smoothing parameter, *τ*, directly lead to increases in the relative mutual information contribution from the prior. Observe that 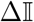 is highly sensitive to the skyline complexity, *p*, thus clarifying how estimates from over-parametrised skyline plots can be dominated by prior information.

Interestingly, we can largely negate the impact of skyline complexity by making *τ* a function of *p*. In the main text we explained how the Skyride implicitly implements the scaling *τ* → *τ*/*p*. While this reduces some of the effect of *p* shown in Fig. 3, it still leads to decaying curves that can, for a given *τ*, be deceptively dependent on smoothing. Here we propose the key transformation *τ* → *τ*/2*p*(*p*–1), as a means of reducing our smoothing in line with our skyline complexity. This transformation was inspired by the dependence of a lower bound on Ω^2^, which we derive in Eq. (A3) later in the Appendix. Its striking impact on the spread of curves from Fig. 3 is given in Fig. A3.

**Fig. A1:**
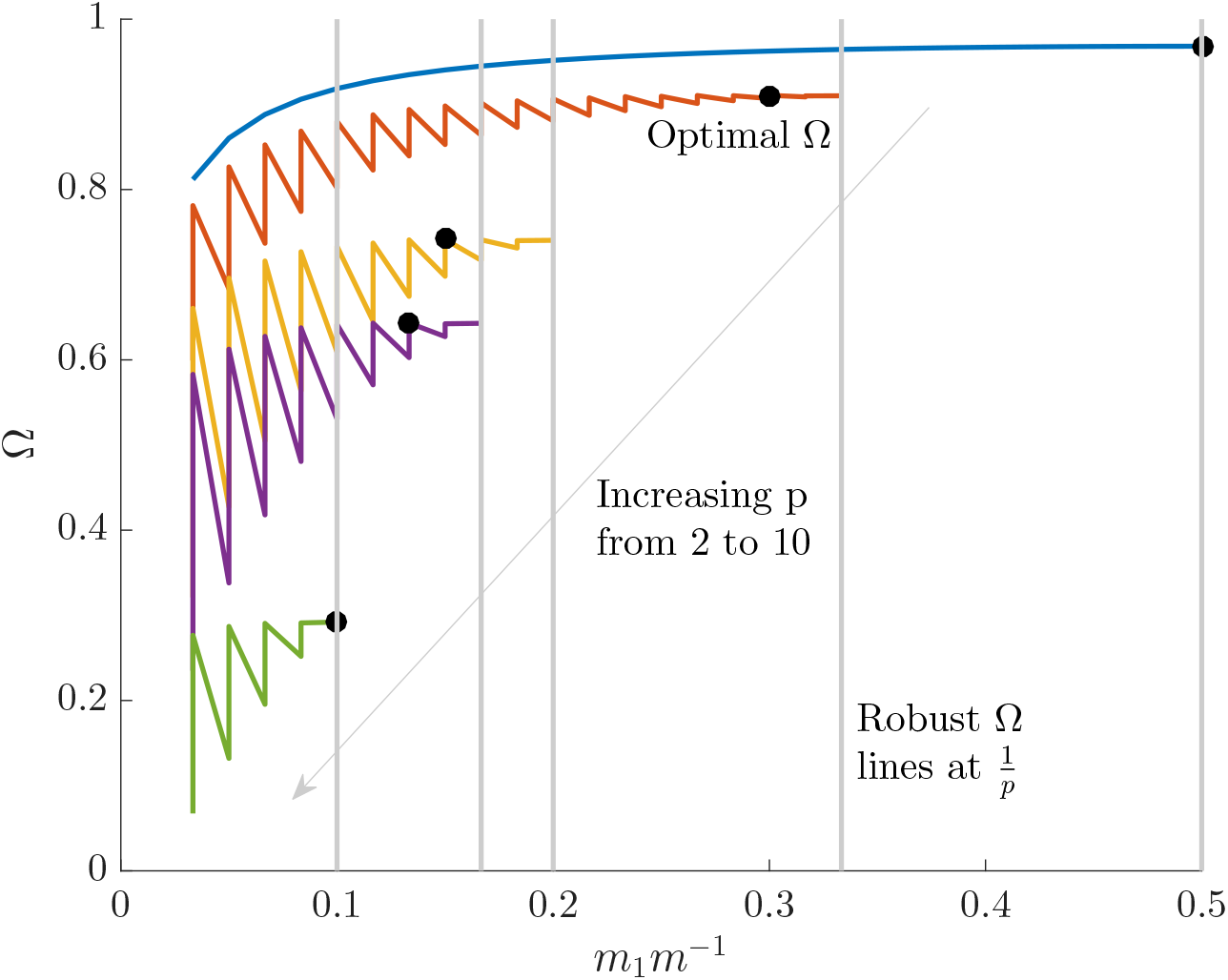
Robust and Ω optimal designs. For the GMRF smoothing prior with *δ_j_* = 1 for all *j* and *τ* = 1, we show that the optimal Ω design point is not always the same as the robust design point, at which 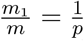. The coloured Ω curves are (along the dashed arrow) for *p* = [2, 3, 5, 6, 10] at *m* = 60, and computed across all partitions for any given *m*_1_ (hence the zig-zagged form). The grey vertical lines mark the robust point for each Ω curve, and the black circles give the optimal Ω points. While these lines and circles do not always match, both generally feature approximately the same Ω values. We found this to be the case across several *m* and *τ* values.

### C. Further Model Selection Bounds

In the the main text we derived lower bounds on Ω^2^, which led to the model rejection parameter, *p*\* (see Eq. (14)). Here we extend and support those results. In Fig. A4 we first show that the bound of Eq. (14) is a good measure of the true Ω^2^ value, for a skyline with uniform GMRF smoothing. We used this bound to define a maximum *p*, *p*\*, above which the skyline would be over-parametrised and susceptible to prior induced overconfidence. We explore *p*\* over *τ* and *m* for this GMRF in Fig. A5 and observe that *p*\* becomes more restrictive with fewer observed data (coalescent events) or increased smoothing. This supports Ω as a useful measure of prior-data contribution.

Lower bounds on Ω^2^ imply upper bounds on the excess mutual information, 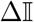 (see Eq. (7)). We manipulate Eq. (14) (under a robust design) to obtain the first inequality in Eq. (A2), with 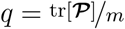 as follows

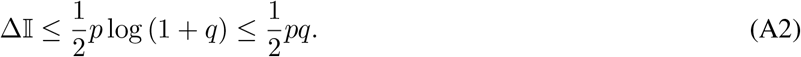

This expression reveals that *p* is akin to a signal bandwidth (by comparison with standard Shannon-Hartley theory [6]) and is therefore a key controlling factor in defining how much additional information the prior will introduce. This supports our proposed *p*\* rejection criterion.

**Fig. A2:**
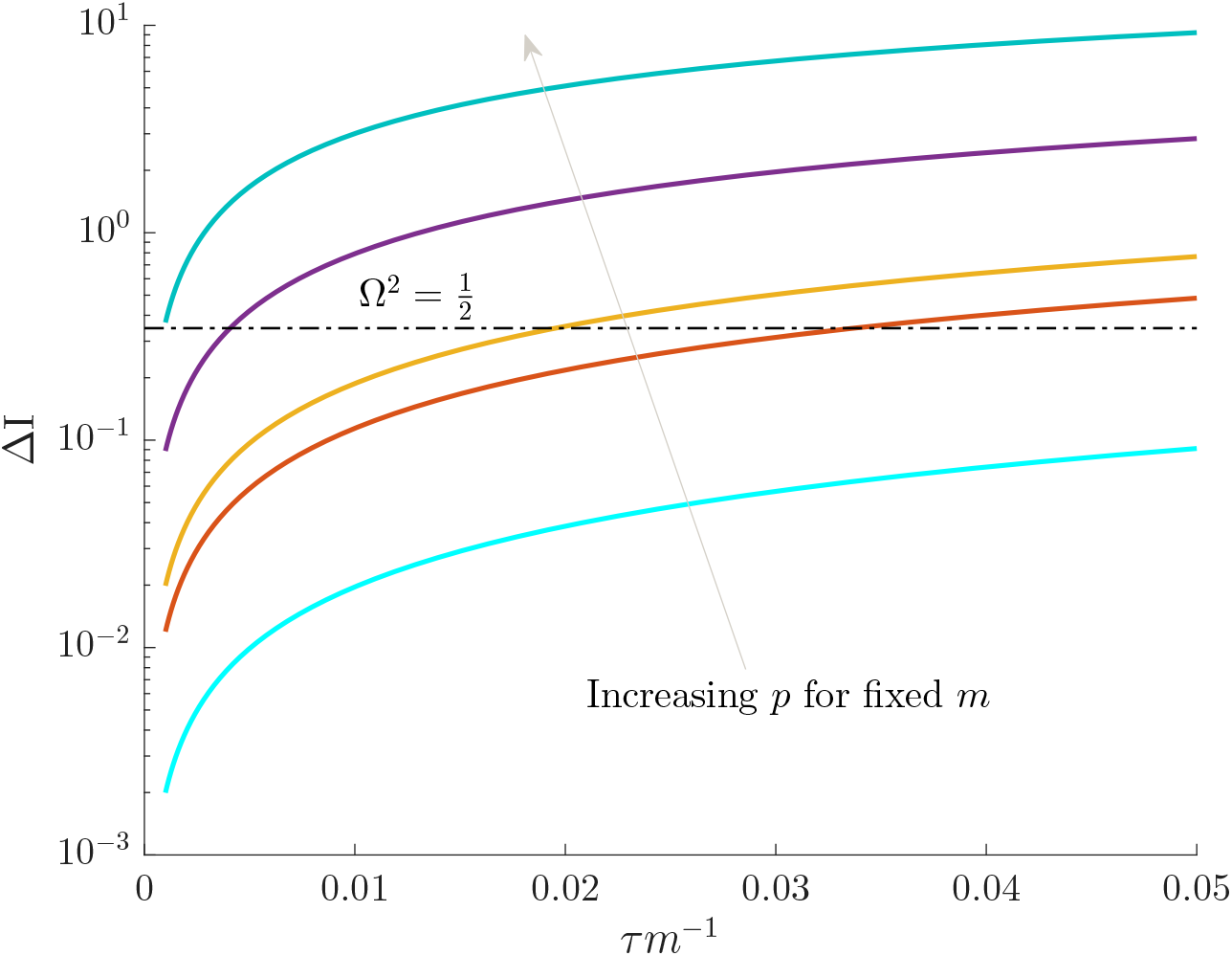
Prior mutual information increases with skyline complexity. For the uniform GMRF, we show that under fixed smoothing (and hence *τ*/*m*), the additional mutual information introduced by the prior, 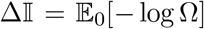, significantly increases with the complexity, *p*, of our skyline. The coloured Ω curves are (along the grey arrow) for *p* = [2, 4, 5, 10, 20] at *m* = 100 with *m_j_* = *m*/*p* (robust design point). The dashed Ω^2^ = 1/2 threshold is also given for comparison. Clearly, the more skyline segments we have for a given tree, the more likely we are being overly informed by our prior.

**Fig. A3:**
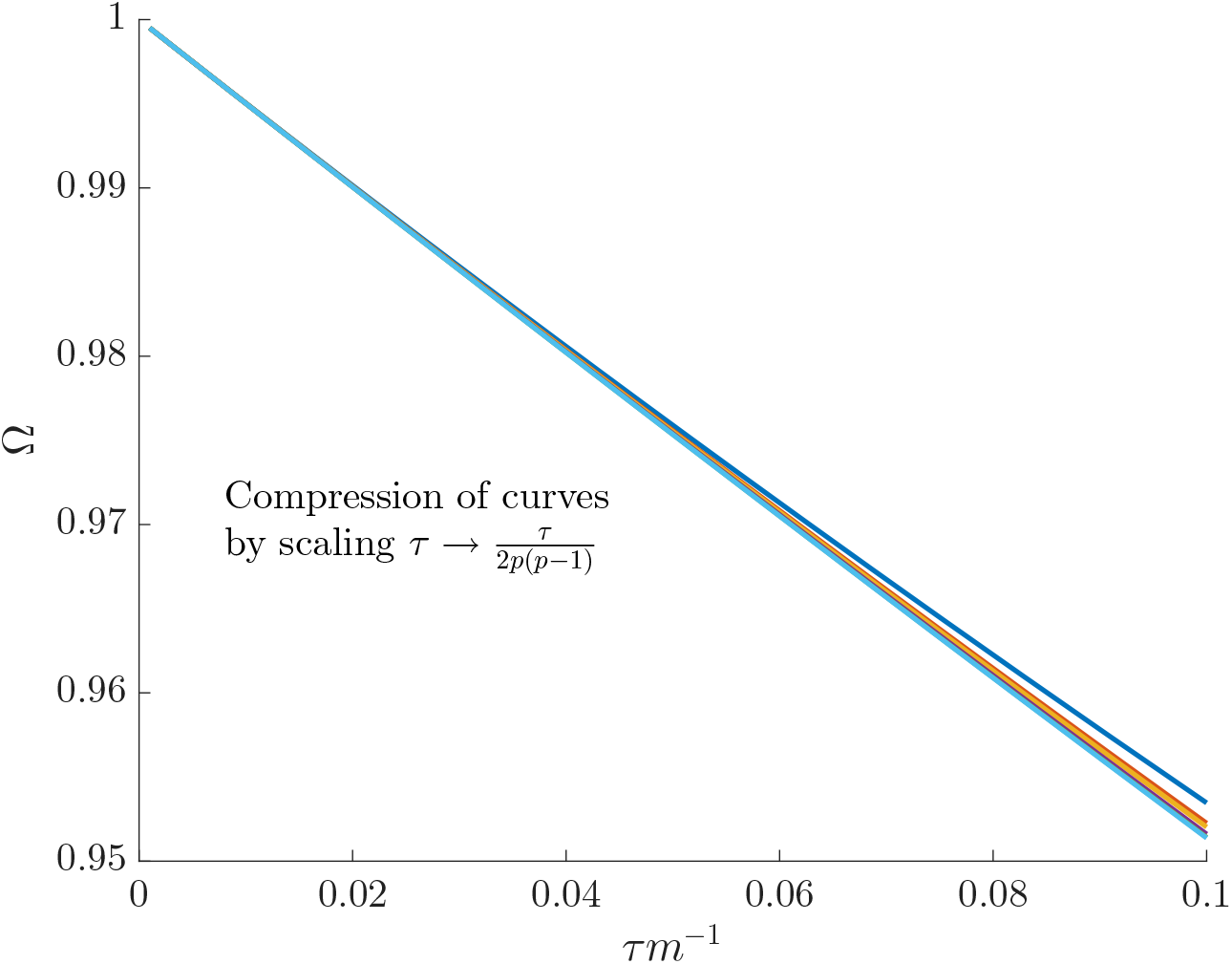
Negating the impact of skyline dimension. We show how an appropriate quadratic scaling of the GMRF precision parameter, *τ*, can remove the complexity (*p*) induced smoothing contribution portrayed in Fig. 3 of the main text. This scaling significantly compresses the coloured Ω curves shown, which are for *p* = [2, 4, 5, 10, 20] at *m* = 100 with *m_j_* = *m*/*p* (robust design point). The resulting Ω^2^ values are now all comfortably above the 1/2 threshold and justified by our information theoretic metrics.

**Fig. A4:**
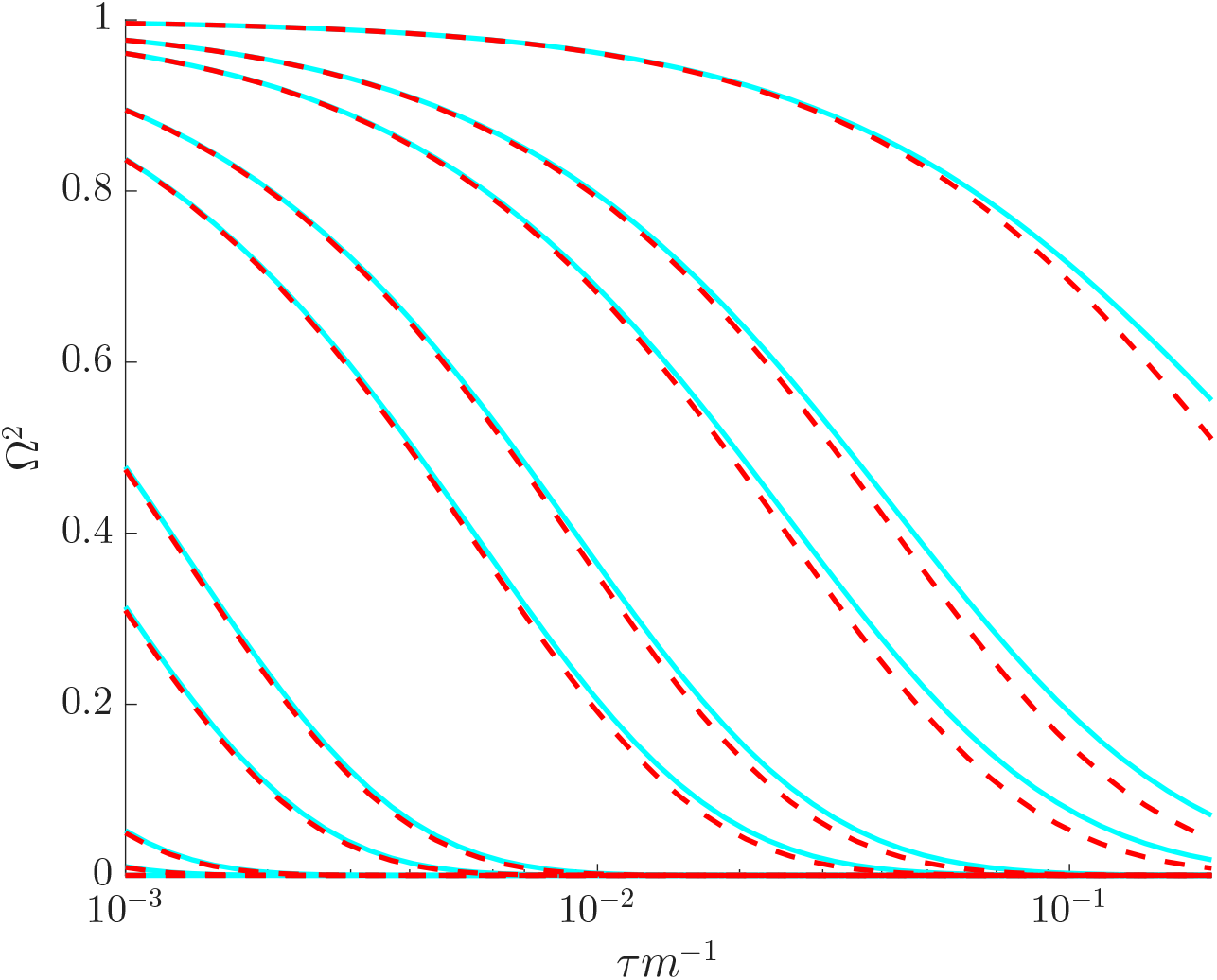
Lower bounds on Ω^2^. For the GMRF smoothing prior with *δ_j_* = 1 for all *j* and *m* = 200, we compare the lower bound on Ω^2^ (red, dashed, see Eq. (14)) with the actual value of Ω^2^ (cyan) at the robust design point of *m_j_* = *m*/*p*. We examine all integer *p* values that are factors of *m*, and find that qualitatively similar comparisons hold for different *τ* and *m* settings. In general the lower bound (*ω*^2^) is a good approximation to Ω^2^.

**Fig. A5:**
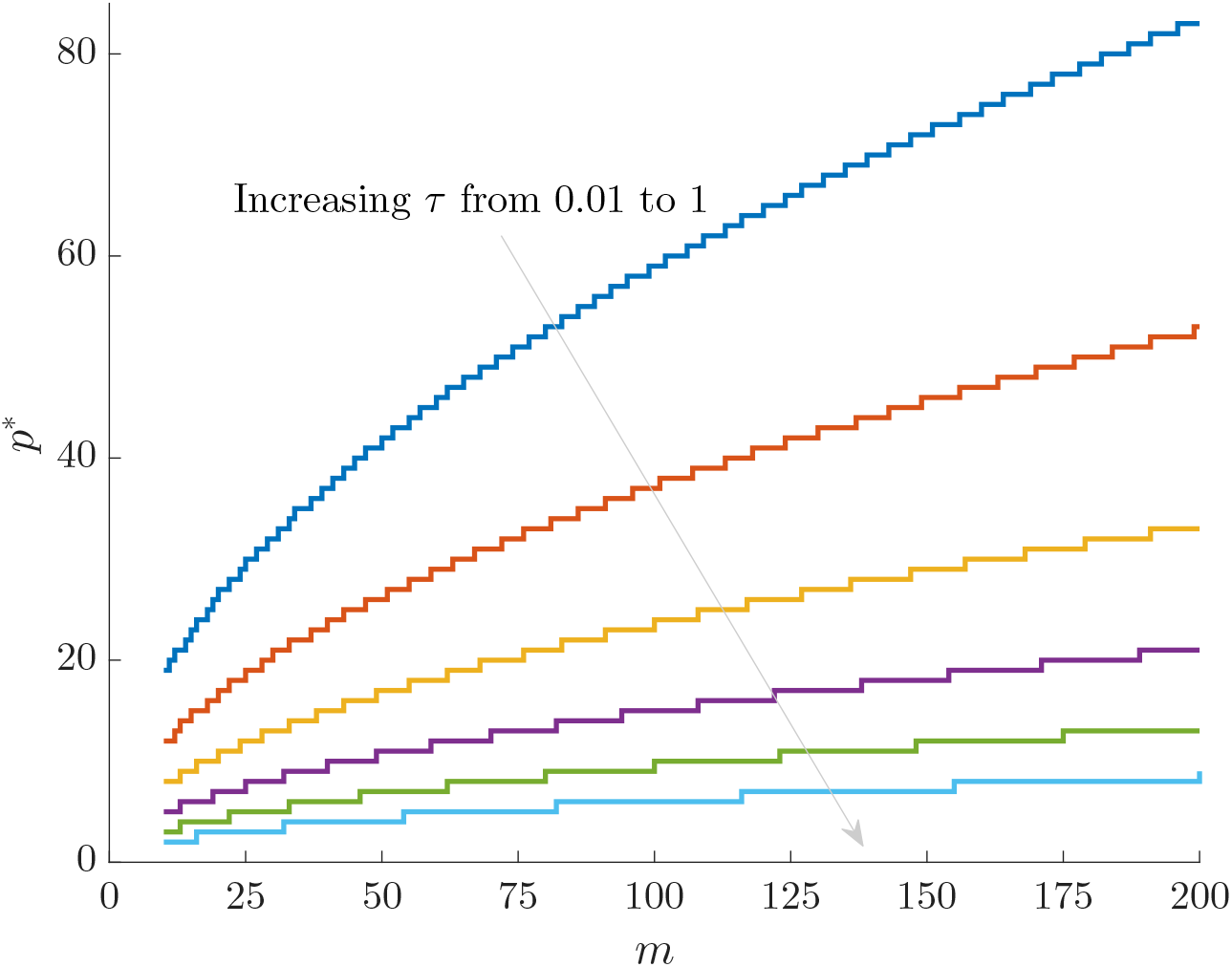
Maximum *p* model selection boundary. For the GMRF smoothing prior with *δ_j_* = 1 for all *j* and at the robust point *m_j_* = *m*/*p*, we compute the maximum allowed number of skyline segments, *p*\*, such that Ω^2^ ≥ 1/2. These curves increase with *m* and decrease with *τ*, indicating how the prior-data contribution can be used to define model rejection regions. Skylines with *p* > *p*\* would be overly informed by the prior and hence should not be used.

Under the log ***N*** parametrisation, 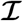 and 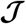 are symmetric, positive definite matrices. For such matrices we can apply a theorem from [14], which states that 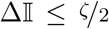, with 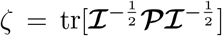. At the robust point, we get 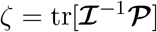, which leads to the second inequality in Eq. (A2). Thus, our bound is tighter than that in [14], and useful for broader, future mathematical analyses of 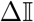. This inequality also clarifies why *m*/*p* is often important for characterising performance here.

We can also use the bound of [14] to derive alternate (but slacker) lower bounds on Ω^2^. This gives the first inequality in Eq. (A3). Applying this to the uniform GMRF gives the second inequality:

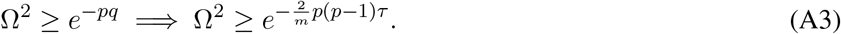

Interestingly, Eq. (A3) shows that the dependence of Ω^2^ on the smoothing parameter *τ* is at most only linear, while the dependence on complexity *p* can be quadratic. This provides further theoretical backing for the use of *p*\* to reject models and emphasises how smoothing can play a deceptively prominent role in the resulting estimate precision produced under complex (high-dimensional) skyline plots.

### D. Ancillary Uncertainty Statistics

In the Egyptian-HCV simulated example we defined two 95% HPD based ancillary statistics for characterising the visual uncertainty present in a skyline plot demographic estimate. In Fig. A6 we plot these statistics and Ω^2^ for various *τ* and *m_j_* values under a time-aware GMRF. We discuss the implications of Fig. A6 in the main text but observe here that trends between the more common (and more easily visualised) HPD based measures and our novel statistic are largely consistent.

**Fig. A6:**
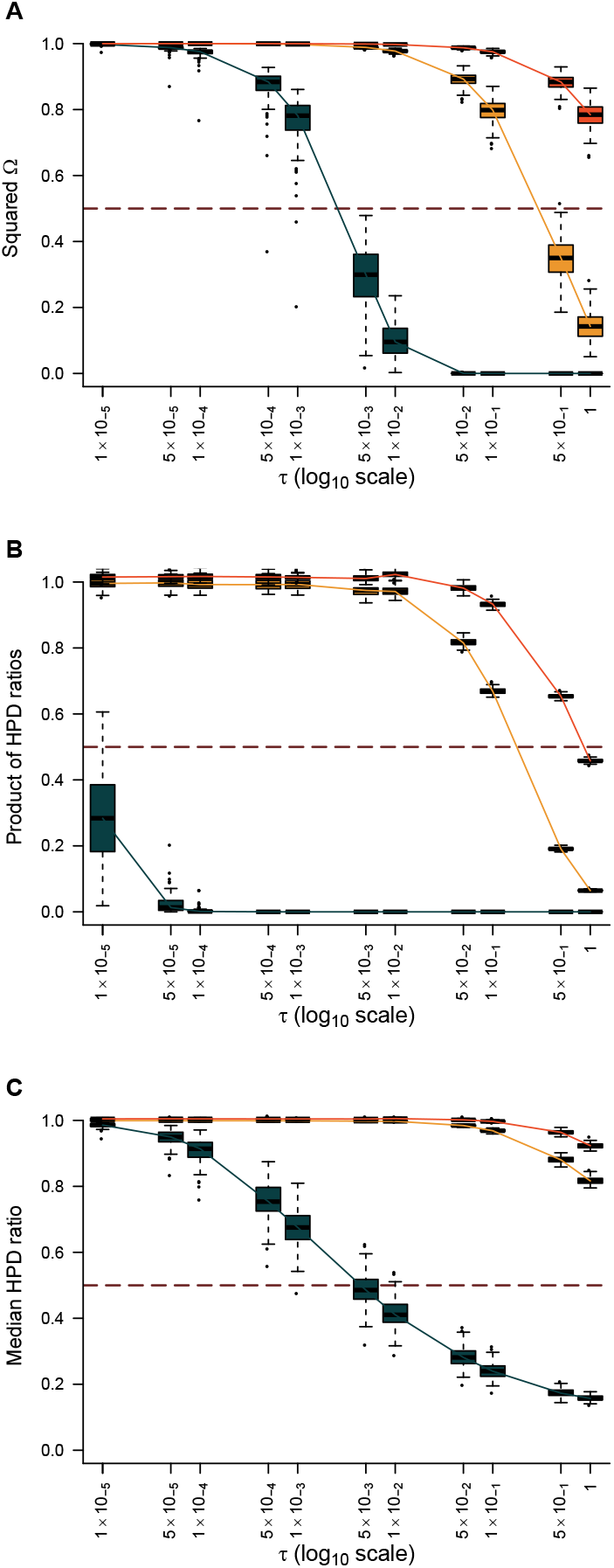
Trends in HPD-based statistics and Ω^2^ under various time-aware GMRF settings. The Ω^2^ (panel A), median HPD ratio of log *N_j_* (panel B) and HPD product (panel C) statistics are computed across log *N_j_* over various combinations of *m_j_* and **τ**. Box-plots summarise our results over 100 observed coalescent trees simulated from previously inferred demographic trends found for the Egyptian HCV dataset. Analyses with *m_j_* = 1 are in dark green, *m_j_* = 4 in yellow and *m_j_* = 8 in orange. The solid lines link the median values across boxes for a given *m_j_* value. The dashed line is positioned at the threshold Ω^2^ = 1/2.

## Notes

### Competing Interest Statement

The authors have declared no competing interest.

### Summary of Updates

Updated text with additional practical example and broader discussion

